# Dna2 nuclease resolves RNA:DNA hybrids at double-strand breaks

**DOI:** 10.1101/2024.08.23.609401

**Authors:** Aditya Mojumdar, Courtney Granger, Jennifer A. Cobb

**Affiliations:** Department of Microbiology & Biochemistry, University of Victoria, Canada

**Keywords:** RNA:DNA hybrids, DSB repair, NHEJ, HR, Dna2 nuclease, Nuclease activity

## Abstract

DNA double-strand breaks (DSBs) are highly detrimental to cells, as improper repair can result in inheritable genetic rearrangements or cell death. The role of RNA:DNA hybrids (RDHs) in DSB repair remains poorly understood, but their transient accumulation and subsequent resolution are crucial for accurate repair. The absence of the end-joining factor Nej1 at DSBs significantly reduced RDH levels, which was linked to increased activity of the Dna2 nuclease. Dna2 limits the accumulation of hybrids at DSBs, with levels rising in the presence of a nuclease-dead Dna2 and *RNH201* deletion. Dna2 has a heightened preference for resolving RDHs with 5’ RNA overhangs compared to duplex substrates with 5’ DNA overhangs. This selective resolution by Dna2 helps restrict hybrid accumulation at DSBs and promotes resection, a function not shared by Exo1. This study underscores the multifunctional roles of canonical repair factors in ensuring efficient homology-directed repair.

**Graphical Abstract:** 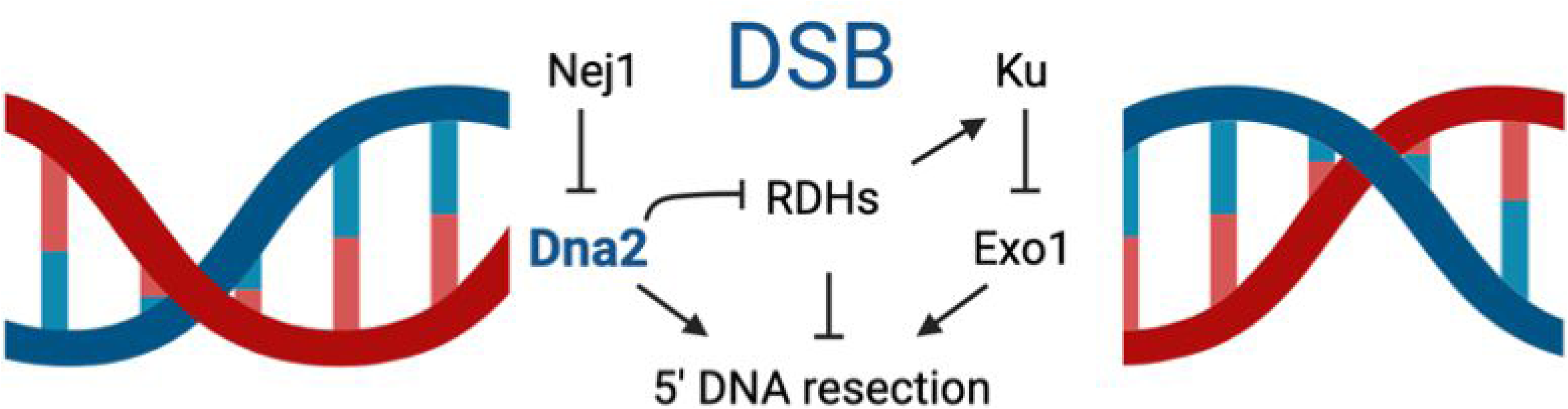

## INTRODUCTION

Double-strand breaks (DSBs) represent a major threat to cellular integrity, as improper repair can result in heritable genetic rearrangements. Cells rely on two primary repair pathways for DSBs: homologous recombination (HR) and non-homologous end joining (NHEJ). HR is an error-free process where DSB ends undergo nucleolytic resection to produce 3’ single-stranded DNA (ssDNA). This ssDNA, along with repair factors, facilitates the invasion of a homologous template. In contrast, NHEJ is an error-prone pathway that involves minimal processing and direct ligation of the DNA ends at the DSB. The regulation of nucleolytic resection plays a critical role in determining which repair pathway is used [1].

Upon the occurrence of a DSB, the Ku70-80 (Ku) complex binds to the DNA ends, primarily to shield them from nucleolytic degradation and to serve as a scaffold for NHEJ-associated proteins [2, 3]. The Mre11-Rad50-Xrs2 (MRX) complex binds to and tethers the loose DSB ends, mainly through Rad50 [4, 5]. Mre11, activated by Sae2, overcomes the Ku barrier and initiates 5’ DNA resection [6, 7]. The MRX complex exhibits exonuclease activity that facilitates short-range resection towards the DSB, while Exo1 and Dna2-Sgs1 show functional redundancy in long-range resection [7–9]. However, Dna2 and Exo1 are regulated differently and have distinct functions at DSBs. In the absence of Mre11, resection primarily occurs through Dna2, independent of Ku. In contrast, Ku binding directly inhibits Exo1 [10–12]. Conversely, Nej1 localization at the DSB inhibits Dna2 but not Exo1 [13–15].

Recent research has revealed that RNA hybrids (RDHs) transiently accumulate at DNA double-strand breaks (DSBs). These hybrids are formed through interactions between DNA at the break site and complementary RNA, which may either be pre-existing or newly transcribed [16, 17]. Understanding the impact of these hybrids on the repair process has proven challenging, as the exact mechanisms governing their formation and removal are still under investigation. Nonetheless, RDHs appear to facilitate both DSB repair and signaling [18–22].

RDHs influence the recruitment of various repair factors and affect DSB repair through both homologous recombination (HR) and non-homologous end joining (NHEJ) [22–28]. Additionally, RDHs have been implicated in RNA-templated DNA repair and may impact the accuracy of DSB repair via NHEJ [29, 30]. While the role of hybrids may vary depending on the stage of repair or specific conditions, they ultimately need to be degraded for repair to proceed successfully. RNases, including type H RNases, and RNA-DNA helicases like Sen1, are known to resolve these hybrids at breaks [18, 25–27, 31]. Given the connections between many DSB repair factors and RNA, and the still poorly defined mechanisms regulating hybrid formation [32], it is crucial to comprehensively investigate the interactions between canonical DSB repair factors and hybrid metabolism. Prior findings indicate that unresolved hybrids impair 5’ resection, a critical step in determining the repair pathway. While hybrids could potentially disrupt 5’ resection directly, increased hybrid accumulation at DSBs has been associated with elevated levels of Ku, suggesting that enhanced end protection by NHEJ factors might reduce 5’ resection [31, 33]. These observations imply that hybrids may form before resection and be removed prior to its initiation, but they do not clarify whether additional repair factors are involved.

In this study, we explore the interplay between RDHs and repair factors in determining repair pathway choice by examining 5’ resection at the HO-DSB. We found that inhibiting hybrid resolution through the deletion of *RNH201* led to an increase in Ku70 accumulation. Conversely, deletion of *NEJ1*, but not *KU70*, resulted in decreased hybrid accumulation, which we attribute to the action of Dna2 nuclease. The antagonistic relationship between Nej1 and Dna2 directly affects RDH metabolism and repair pathway choice. Our findings show that Dna2 resolves hybrids, and RDH levels increase significantly in *dna2*-1 mutants lacking nuclease activity. While unresolved hybrids were previously thought to impair 5’ resection, it was unclear whether this effect was direct or mediated by increased Ku levels [31, 33]. Our results demonstrate that while Ku70-mediated end protection was a key factor, 5’ resection and hybrid levels can be mechanistically uncoupled. These findings offer valuable insights into the complex interactions between RNA:DNA hybrids, Dna2, and other DNA double-strand break (DSB) repair factors. The crucial role of Dna2 in regulating RDH accumulation highlights its essential function in maintaining genomic stability during DSB repair. Furthermore, the initiation of hybrid resolution by Dna2 sheds light on the additional mechanisms that cells utilize to ensure effective homology-directed repair.

## RESULTS

### Hybrids Regulate Resection at Double-Strand Breaks

To elucidate the role of RNA hybrids (RDHs) in double-strand break (DSB) repair, we utilized the inducible HO endonuclease system in budding yeast (Figure 1A). This system, well-established for studying DSB repair in eukaryotes, creates a single DNA DSB at the *MAT* locus upon galactose induction [34]. Our goal was to determine how RDHs interact with DSB repair factors to influence repair pathway choice.

**Figure 1.**
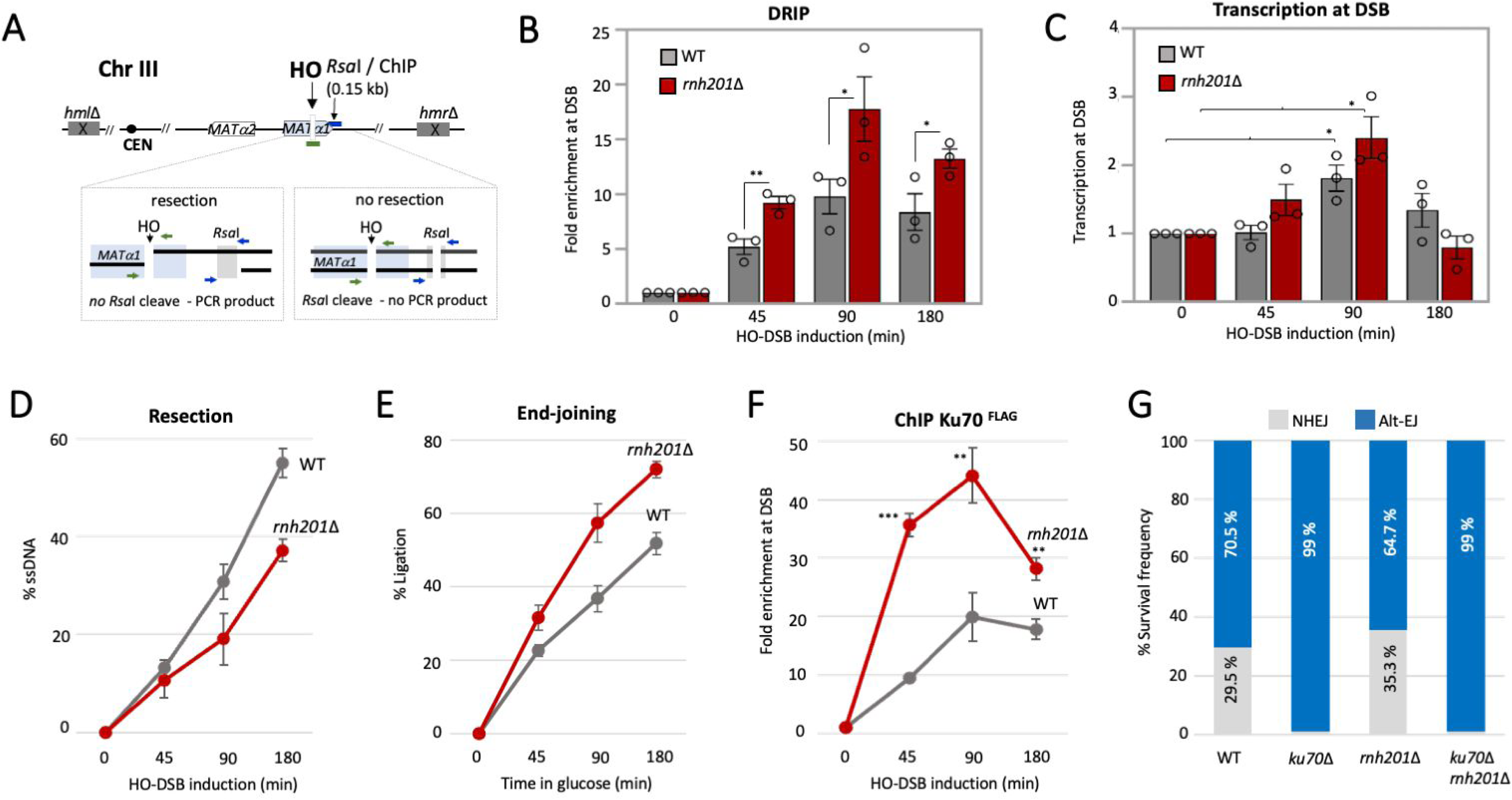
Hybrids Regulate Resection at DSB (A) Schematic representation of regions around the HO cut site on chromosome III. The ChIP primers used in this study correspond to 0.15kb from the DSB and the end-joining primers flank the HO cut site (Lig). The qPCR resection assay relies on two RsaI sites located 0.15 kb from the DSB. **(B)** Enrichment of RNA:DNA hybrids (RDHs) using S9.6 antibody at 0.15kb from DSB, at 0, 45, 90 and 180 mins post DSB induction in cycling cells in wild type (JC-727), and *rnh201*Δ (JC-5614) was determined. The fold enrichment is normalized to recovery at the *PRE1* locus. **(C)** Transcription at 0.15kb from DSB, at 45, 90 and 180 mins post DSB induction in cycling cells relative to 0 min (no DSB induction), in wild type (JC-727), and *rnh201*Δ (JC-5614) was determined and normalized to the *ACT1* locus. **(D)** qPCR-based resection assay of DNA 0.15 kb away from the HO-DSB, as measured by % ssDNA, at 0, 45, 90 and 180 mins post DSB induction in cycling cells in wild type (JC-727) and *rnh201*Δ (JC-5614). **(E)** qPCR-based ligation assay of DNA at the HO-DSB, as measured by % ligation. Cells were induced with galactose for 3 hours before wash and release into glucose at 0, 45, 90 and 180 mins in wild type (JC-727), and *rnh201*Δ (JC-5614). **(F)** Enrichment of Ku70^Flag^ at 0.15kb from DSB, at 0, 45, 90 and 180 mins post DSB induction in cycling cells in wild type (JC-3964), and *rnh201*Δ (JC-5950) was determined. The fold enrichment is normalized to recovery at the *PRE1* locus. **(G)** Survival frequencies depicting the ratio of NHEJ (gray) and Alt-EJ/MMEJ (blue) repair frequencies in wild type (JC-5903), *ku70*Δ (JC-6195), *rnh201*Δ (JC-6198), and *ku70*Δ *rnh201*Δ (JC-6288). For all the experiments -error bars represent the standard error of three replicates. Significance was determined using a 1-tailed, unpaired Student’s t test (*p* < 0.05∗; *p* < 0.01∗∗; *p* < 0.001∗∗∗). All strains compared are marked and compared to WT.

Initially, we performed DNA-RNA immunoprecipitation (DRIP) using the S9.6 antibody to quantify hybrid levels at the DSB. Consistent with previous studies, hybrids were detected as early as 45 minutes post-HO induction and peaked at 90 minutes (Figure 1B) [31, 33]. RDH accumulation was notably higher in *rnh201*Δ mutants, lacking RNaseH2 which degrades RNA in RDHs (Figure 1B) [18, 35]. This result differed from observations in cells blocked in G2/M phase with nocodazole prior to HO cutting [31]. Early hybrid formation at 45 minutes likely originates from pre-existing RNA, as the HO cut site is located within an intragenic region (Figure 1C). By 90 minutes, increased RDHs could result from both pre-existing RNA and slight transcriptional increases. However, the elevated RDH levels in *rnh201*Δ were attributed to reduced hybrid resolution post-formation, with no significant difference in transcription levels between wild type and *rnh201*Δ cells at the various timepoints. Hybrids decreased with *in vitro* RNaseH1 treatment or *in vivo* RNaseH1 overexpression (Figure S1A).

We next examined whether increased hybrid levels altered repair pathway choices by assessing resection, a critical step that directs the pathway of repair. Resection was measured using a method involving an RsaI cut site 0.15 kb from the DSB (Figure 1A) [36, 37]. If resection extends beyond this recognition site, single-stranded DNA is produced, preventing RsaI cleavage and leading to locus amplification by PCR. Resection levels decreased over time in *rnh201*Δ mutants compared to wild type (Figure 1D). Given the transient nature of hybrids and their potential accumulation at various stages of repair, reduced resolution could impact repair outcomes. We assessed end-ligation post-HO cutting, when cleavage was >90%, by quantitative PCR with primers flanking the DSB [14]. In *rnh201*Δ mutants, end-joining capacity was increased compared to wild type (Figure 1E).

To determine whether increased end-joining correlated with altered recovery of end-joining factors, we performed ChIP at the DSB. Ku70^FLAG^ levels rose early after DSB formation and remained elevated in *rnh201*Δ mutants compared to wild type (Figure 1F). Ku70^FLAG^ recovery peaked at 90 minutes, coinciding with the peak of RDH accumulation (Figures 1B and 1F) [31, 33]. By contrast, other end-joining factors like Nej1, Lif1 and Dnl4, showed similar enrichment in wild type and *rnh201Δ* mutants (Figure S1B-D). We then assessed the frequency of NHEJ and MMEJ in relation to end-ligation and hybrid levels using a reporter system with a *URA3* marker flanked by inverted HO recognition sites (Figure S1E) [38]. Simultaneous cleavage of both sites generates non-compatible ends, leading to alt-EJ/MMEJ, but single cuts can still be repaired by NHEJ [15, 38]. We observed a slight increase in NHEJ relative to MMEJ in *rnh201*Δ mutants, aligning with increased Ku presence at the break (Figure 1G). End-ligation increased and resection decreased when hybrid levels increased, warranting further investigation into the interplay between DSB repair factors and RDHs in the regulation of 5’ resection.

### Deletion of *NEJ1* Reduces RDH Accumulation

Having established that increased RDHs correlate with enhanced Ku and NHEJ, and decreased end-resection, we investigated whether end-joining factors influence RDH levels. DRIP was performed in *ku70*Δ and *nej1*Δ mutant cells. Ku70 and Nej1 are essential for NHEJ and inhibit Exo1 and Dna2, respectively, which drive resection at the DSB [13–15, 37, 39]. RDH levels were similar in *ku70*Δ and wild type, with no significant difference in hybrid levels between *rnh201*Δ and *ku70*Δ *rnh201*Δ mutants (Figure 2A and 2B). The double mutant allowed us to discern whether hybrid accumulation impacts resection directly or through increased Ku. Early resection levels were similarly reduced in *rnh201*Δ and *ku70*Δ *rnh201*Δ mutants up to 90 minutes post-HO cutting, suggesting direct interference by RDHs in resection, independent of Ku status (Figure 2C). At the 180-minute timepoint, resection increased to slightly above wild type if *KU70* was deleted in *rnh201*Δ, indicating that Ku70 influences resection when hybrid levels are high after 90 minutes, but not at early timepoints. DRIP at corresponding timepoints revealed similar hybrid levels in *rnh201*Δ and *ku70*Δ *rnh201*Δ mutants (Figure 2B), suggesting that at later timepoints, RDHs do not directly interfere with resection but may do so indirectly via increased Ku. In *ku70*Δ *rnh201*Δ mutants, resection remained slow from 0-90 minutes but increased rapidly from 90-180 minutes (Figure 2C).

**Figure 2.**
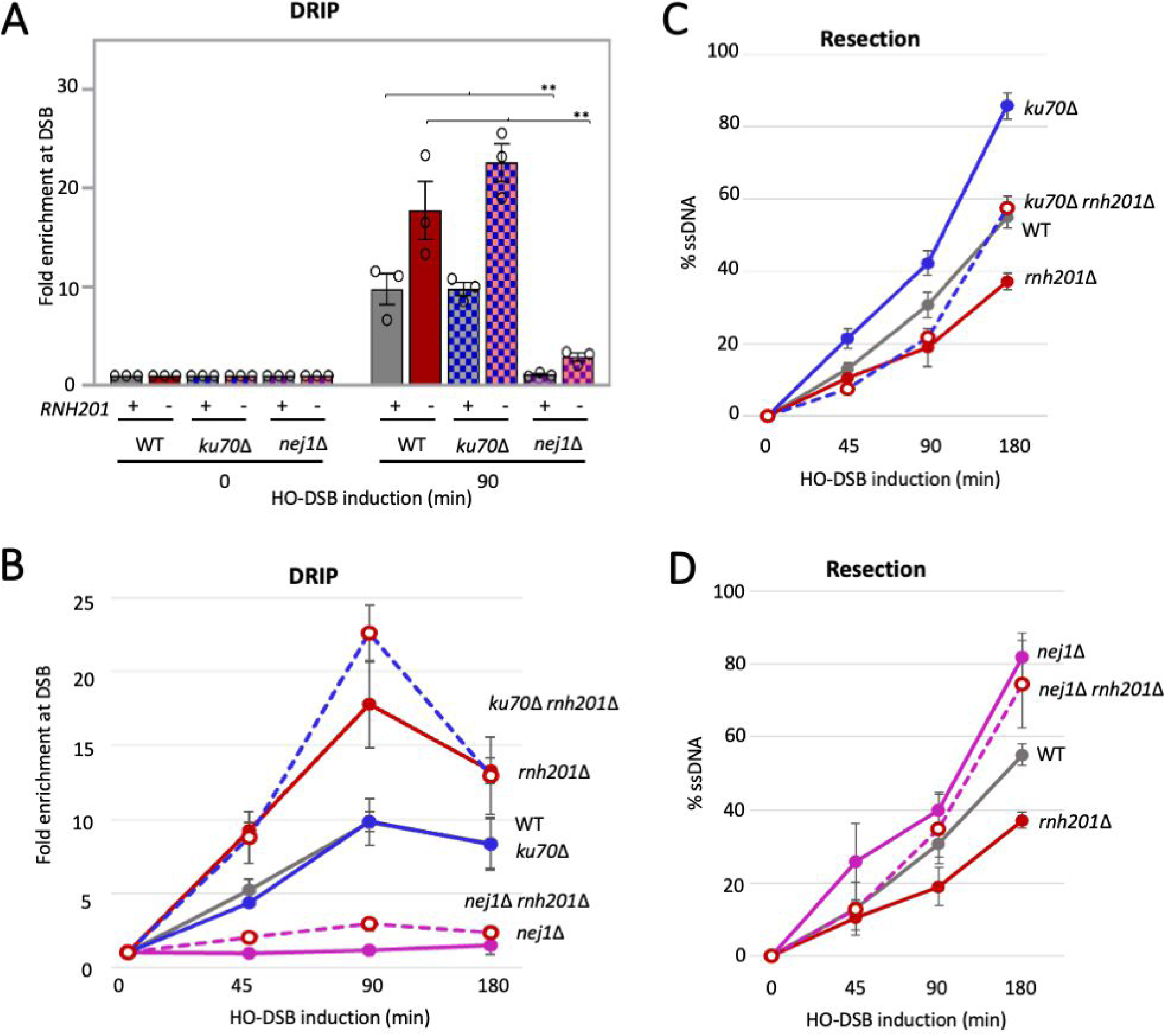
Deletion of NEJ1 Reduces RDH Accumulation (A) Enrichment of RDHs using S9.6 antibody at 0.15kb from DSB, 90 mins after DSB induction relative to 0 min (no DSB induction), in wild type (JC-727), *rnh201*Δ (JC-5614), *ku70*Δ (JC-1904), *ku70*Δ *rnh201*Δ (JC-5720), *nej1*Δ (JC-1342), and *nej1*Δ *rnh201*Δ (JC-5756) was determined. The fold enrichment is normalized to recovery at the *PRE1* locus. **(B)** Enrichment of RDHs using S9.6 antibody at 0.15kb from DSB, at 0, 45, 90 and 180 mins post DSB induction in cycling cells in wild type (JC-727), *rnh201*Δ (JC-5614), *ku70*Δ (JC-1904), *ku70*Δ *rnh201*Δ (JC-5720), *nej1*Δ (JC-1342), and *nej1*Δ *rnh201*Δ (JC-5756) was determined. The fold enrichment is normalized to recovery at the *PRE1* locus. **(C)** qPCR-based resection assay of DNA 0.15 kb away from the HO-DSB, as measured by % ssDNA, at 0, 45, 90 and 180 mins post DSB induction in cycling cells in wild type (JC-727), *rnh201*Δ (JC-5614), *ku70*Δ (JC-1904), and *ku70*Δ *rnh201*Δ (JC-5720). **(D)** qPCR-based resection assay of DNA 0.15 kb away from the HO-DSB, as measured by % ssDNA, at 0, 45, 90 and 180 mins post DSB induction in cycling cells in wild type (JC-727), *rnh201*Δ (JC-5614), *nej1*Δ (JC-1342), and *nej1*Δ *rnh201*Δ (JC-5756). For all the experiments - error bars represent the standard error of three replicates. Significance was determined using a 1-tailed, unpaired Student’s t test (*p* < 0.05∗; *p* < 0.01∗∗; *p* < 0.001∗∗∗). All strains compared are marked and compared to WT.

A marked decrease in RDHs was observed in *nej1*Δ mutants, contrasting with *ku70*Δ (Figure 2A and 2B). Neither *ku70*Δ nor *nej1*Δ mutants showed altered transcription at DSBs, indicating that reduced RDH levels in *nej1*Δ mutants result from increased hybrid resolution (Figure S1F). Notably, *NEJ1* deletion reversed the increased RDH levels in *rnh201*Δ to below wild type levels, similar to *nej1*Δ single mutants (Figure 2A and 2B). This suggests that Nej1 loss at DSBs enhances hybrid resolution through a mechanism distinct from RNaseH1 regulation. Furthermore, from 0-90 minutes post-HO cutting, resection in *nej1*Δ *rnh201*Δ mutants increased similarly to nej1Δ (Figure 2D), indicating that Nej1 presence negatively regulates resection, potentially through its impact on RDHs at early timepoints.

### Dna2 Modulates RDH Accumulation at Double-Strand Breaks

To further understand the regulation of RNA-DNA hybrids (RDHs) at double-strand breaks (DSBs), we explored the effects of Dna2 in *nej1*Δ mutants, which show reduced RDH levels. Dna2 is known for processing small RDHs, such as those from Okazaki fragments during DNA replication [40–44]. Our previous research indicated that Nej1 downregulates Dna2 recovery at DSBs [37]. In this study, we observed a significant increase in Dna2^HA^ localization at DSBs in *nej1*Δ mutants 45 minutes after galactose induction (Figure 3A). This suggests that Dna2 recruitment to DSBs is enhanced early after break formation and that its retention is prolonged in the absence of Nej1. These data also highlight the antagonistic relationship of Dna2 and Nej1 early after DSB formation when repair pathway choice is being determined and suggests the interplay between these repair factors directly impacts RDH metabolism at DSBs.

**Figure 3.**
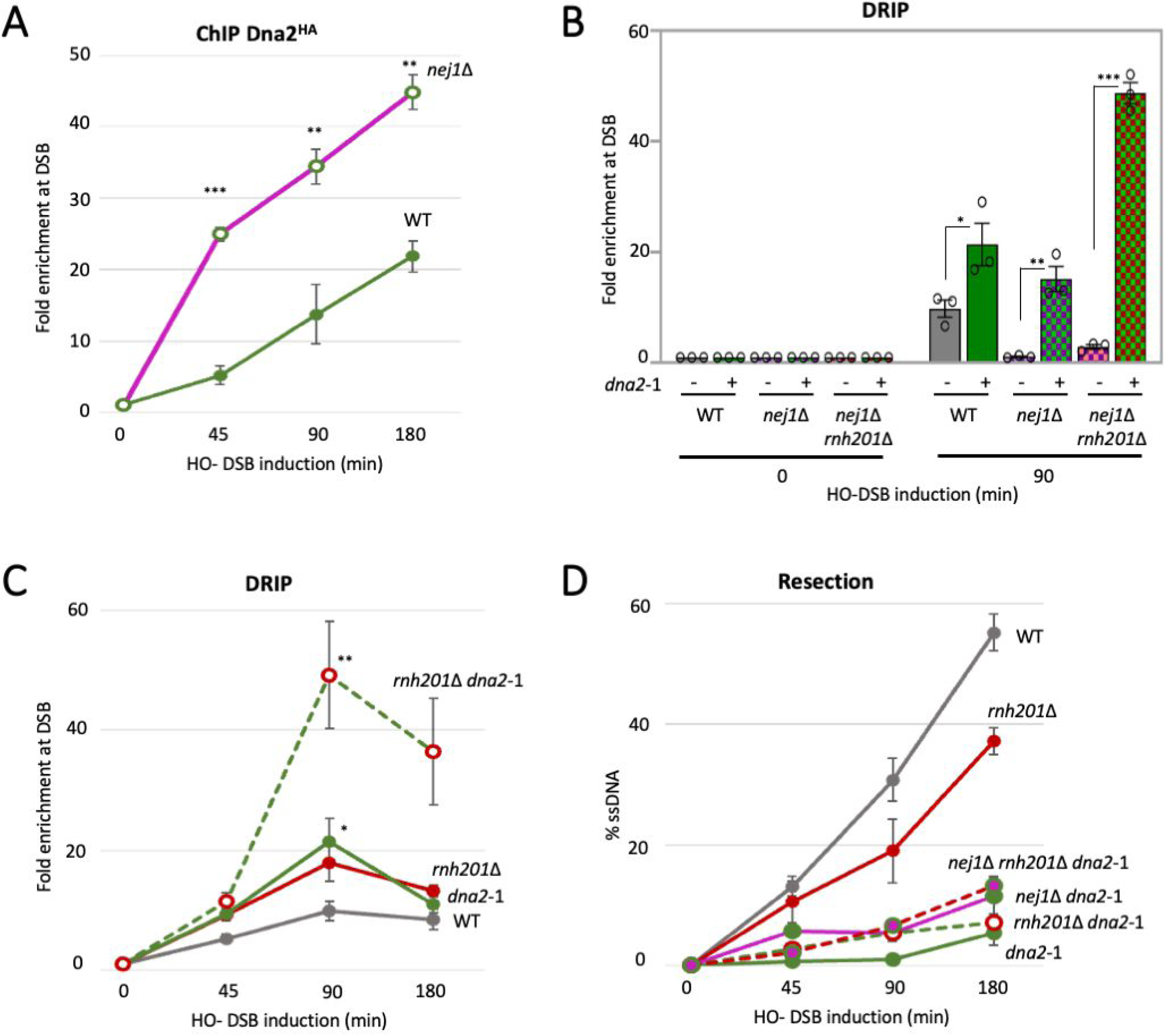
Dna2 Modulates RDH Accumulation at Double-Strand Breaks (A) Enrichment of Dna2^HA^ at 0.15kb from DSB, at 0, 45, 90 and 180 mins post DSB induction in cycling cells in wild type (JC-4117) and *nej1*Δ (JC-4118) was determined. The fold enrichment is normalized to recovery at the *PRE1* locus. **(B)** Enrichment of RDHs using S9.6 antibody at 0.15kb from DSB, 90 mins after DSB induction relative to 0 min (no DSB induction), in wild type (JC-727), *dna2-*1 (JC-6007), *nej1*Δ (JC-1342), *nej1*Δ *dna2-*1 (JC-5670), *nej1*Δ *rnh201*Δ (JC-5756), and *nej1*Δ *dna2-*1 *rnh201*Δ (JC-5733) was determined. The fold enrichment is normalized to recovery at the *PRE1* locus. **(C)** Enrichment of RDHs using S9.6 antibody at 0.15kb from DSB at 0, 45, 90 and 180 mins post DSB induction in cycling cells in wild type (JC-727), *rnh201*Δ (JC-5614), *dna2-*1 (JC-6007), and *dna2-*1 *rnh201*Δ (JC-5734), was determined. The fold enrichment is normalized to recovery at the *PRE1* locus. **(D)** qPCR-based resection assay of DNA 0.15 kb away from the HO-DSB, as measured by % ssDNA, at 0, 45, 90 and 180 mins post DSB induction in cycling cells in wild type (JC-727), *rnh201*Δ (JC-5614), *dna2-*1 (JC-6007), *dna2-*1 *rnh201*Δ (JC-5734), *nej1*Δ (JC-1342), *nej1*Δ *dna2-*1 (JC-5670), *nej1*Δ *rnh201*Δ (JC-5756), and *nej1*Δ *dna2-*1 *rnh201*Δ (JC-5733). For all the experiments -error bars represent the standard error of three replicates. Significance was determined using a 1-tailed, unpaired Student’s t test (*p* < 0.05∗; *p* < 0.01∗∗; *p* < 0.001∗∗∗). All strains compared are marked and compared to WT.

To investigate this further, we examined RDH levels in cells carrying the *dna2*-1 allele, which harbors a point mutation (P504→S) that abolishes nuclease activity but does not affect Dna2 recruitment to DSBs [15, 41]. When *dna2*-1 was combined with *nej1*Δ and *nej1*Δ *rnh201Δ*, we observed a marked increase in hybrid accumulation 2-fold and 5-fold above wild type, respectively (Figure 3B). This indicates that Dna2 nuclease activity is crucial for reducing RDH levels at DSBs. Additionally, *dna2*-1 exhibited elevated RDH levels, underscoring the essential role of Dna2 in hybrid resolution under normal conditions, not solely when its levels are increased due to *NEJ1* deletion (Figure 3B and 3C).

In *dna2*-1 *rnh201*Δ double mutants, there was a significant additive increase in RDH accumulation (Figure 3B and 3C). The extent of DNA resection in both the double mutant and *dna2*-1 was notably reduced compared to *rnh201*Δ, highlighting direct involvement of Dna2 in 5’ DNA resection (Figure 3D). Our previous research demonstrated that Ku70^FLAG^ levels were elevated in *dna2*-1, and this resection defect was mitigated by deleting *KU70* [15]. Although Ku70^FLAG^ levels also additively increased in *dna2*-1 *rnh201*Δ, deleting *KU70* only modestly restored resection in these double mutants, unlike in *dna2*-1 *ku70*Δ where resection surpassed wild-type levels (Figure S2A-B) [15]. These results suggest that in cells with increased hybrids, elevated Ku70^FLAG^ is not the sole factor contributing to reduced resection.

Importantly, attributing resection defects solely to increased RDH levels in *rnh201*Δ mutants may be an oversimplification, as the recovery of both Exo1^HA^ and Dna2^HA^ at the break decreased by approximately 50% compared to wild type (Figure S3A-B). The complex interplay among these nucleases and their functional redundancy in DNA resection complicates understanding their roles in hybrid resolution. Nevertheless, in contrast to *dna2*-1, RDH levels in *exo1*Δ mutants were comparable to wild type, and *exo1*Δ *rnh201*Δ double mutants did not exhibit significant differences in hybrid accumulation compared to *rnh201*Δ (Figure S2C). This highlights a Rnh201-independent function of Dna2 in hybrid resolution, distinct from Exo1. Of note, recovery of Rad50^HA^, a MRX complex subunit, and the nuclease associated factors, Sae2^HA^ and Sgs1^HA^, were similarly recovered in *rnh201*Δ and wild type cells (Figure S3D-F).

### Dna2 Nuclease in RDH Resolution

To explore whether Dna2 directly resolves RDHs and facilitates DNA resection, we characterized Dna2 *in vitro*, assessing its substrate specificity. We expressed and purified Dna2 from yeast, confirmed its folding through a thermal shift assay (Figure S4A-B), and tested its activity on various DNA and RNA hybrid substrates. As a 5’ to 3’ exonuclease, Dna2 was evaluated using 3’ FAM-labeled substrates (Figure 4A). Our assays revealed that Dna2 hydrolyzed ssDNA and dsDNA with a 5’ overhang similarly (Figure S4C-D) but showed a preference for RDHs with a 5’ RNA overhang (Figure 4B).

**Figure 4.**
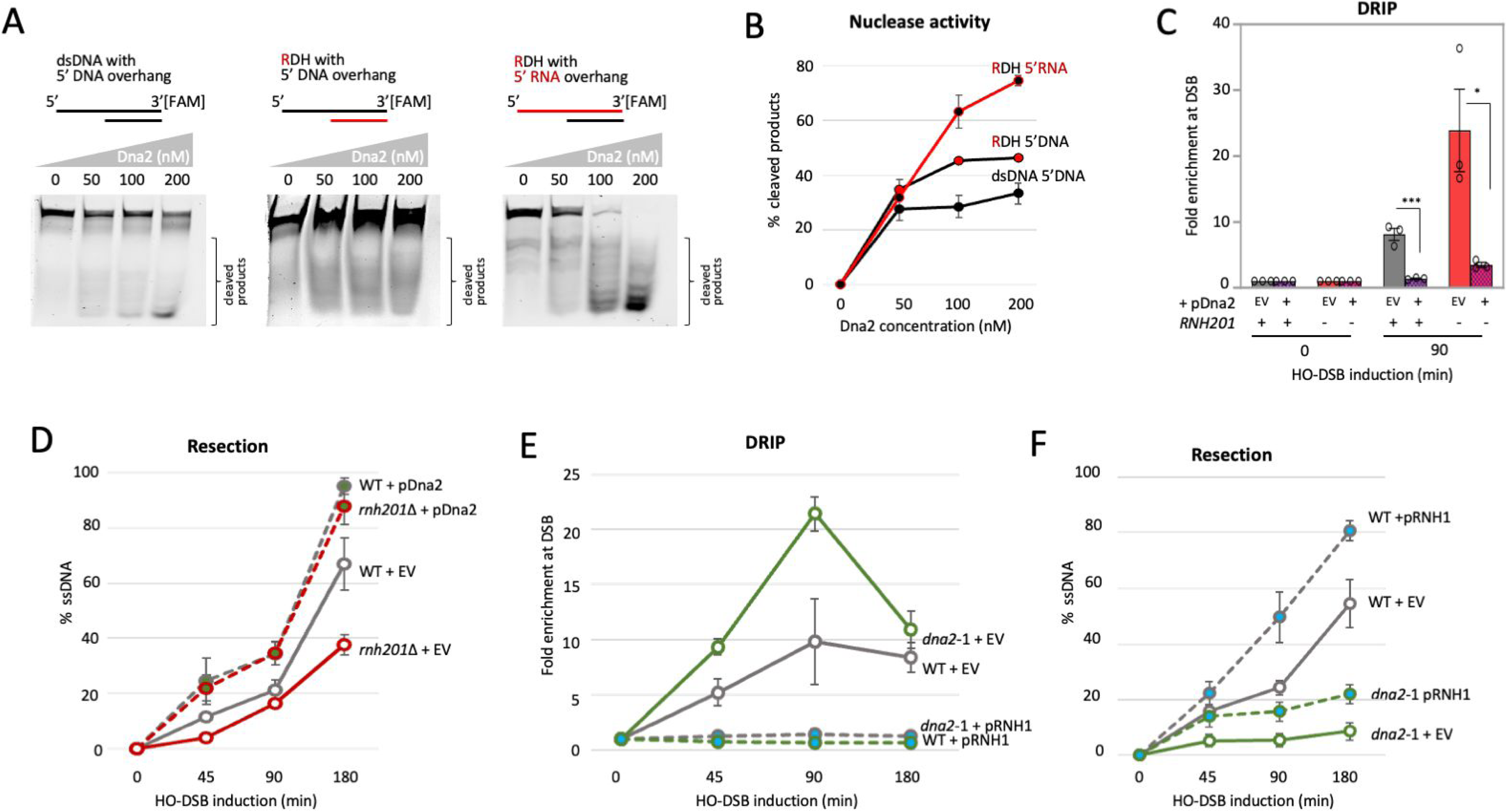
Dna2 Nuclease in RDH Resolution (A) Gel images showing the nuclease activity of purified Dna2 (0 to 200 nM) with 50 nM of substrates – dsDNA, DNA:RNA hybrid with 5’ DNA overhang and RNA:DNA hybrid with 5’ RNA overhang. **(B)** Plot of the quantified bands corresponding to the degradation products shown in (B) normalized to the total substrate and represented as % degradation products. **(C)** Enrichment of RDHs using S9.6 antibody at 0.15kb from DSB, 90 mins after DSB induction relative to 0 min (no DSB induction), in wild type (JC-727), and *rnh201*Δ (JC- 5614), either with empty vector pGAL LexA or with vector expressing Dna2 (pGAL LexA-Dna2) was determined. The fold enrichment is normalized to recovery at the *PRE1* locus. **(D)** qPCR-based resection assay of DNA 0.15 kb away from the HO-DSB, as measured by % ssDNA, at 0, 45, 90 and 180 mins post DSB induction in cycling cells in wild type (JC-727), and *rnh201*Δ (JC-5614), either with empty vector pJG4-7 or with vector expressing Dna2 (pJG4-7-Dna2). **(E)** Enrichment of RNA:DNA hybrids using S9.6 antibody at 0.15kb from DSB at 0, 45, 90 and 180 mins post DSB induction in cycling cells in wild type (JC-727) and *dna2*-1 (JC-6007), either with empty vector pRS425 or with vector expressing RNH1 (pRS425-RNH1) was determined. The fold enrichment is normalized to recovery at the *PRE1* locus. **(F)** qPCR-based resection assay of DNA 0.15 kb away from the HO-DSB, as measured by % ssDNA, at 0, 45, 90 and 180 mins post DSB induction in cycling cells in wild type (JC-727) and *dna2*-1 (JC-6007), either with empty vector pRS425 or with vector expressing RNH1 (pRS425-RNH1). For all the experiments - error bars represent the standard error of three replicates. Significance was determined using a 1-tailed, unpaired Student’s t test (*p* < 0.05∗; *p* < 0.01∗∗; *p* < 0.001∗∗∗). All strains compared are marked and compared to WT.

When Dna2 was overexpressed in *rnh201*Δ mutants, RDH levels significantly decreased, demonstrating that enhanced Dna2 activity can reverse increased hybrid accumulation *in vivo* (Figure 4C). This decrease in RDHs was associated with a corresponding increase in DNA resection above wild type levels (Figure 4D). However, despite increased resection via Exo1 overexpression in *rnh201*Δ, RDH levels did not decrease (Figure S5A-B), suggesting that hybrid resolution precedes DNA resection and is specifically mediated by Dna2, rather than Exo1 at DSBs. Conversely, overexpressing Rnh1 reduced RDH levels in *dna2*-1 mutants to below wild type levels with empty vector controls (Figure 4E). Resection in *dna2*-1 + RNH1 was modestly enhanced compared to *dna2*-1 + EV, though it remained below wild type levels (Figure 4F). This indicates that while Dna2 directly contributes to DNA resection, its effect on RDHs also indirectly influences resection. The partial restoration of resection in *dna2*-1 mutants with increased RDH levels further highlights the complex relationship between hybrid resolution and resection.

## DISCUSSION

Using the HO-DSB system in budding yeast, we demonstrate that RDHs (RNA-DNA hybrids) accumulate at the break site within 45 minutes of DSB (double-strand break) induction, and before significant changes in transcription are detected. Peak hybrid levels are observed around 90 minutes, which is also an inflection point where the impact of hybrids on resection becomes Ku dependent. While the literature is divided on whether RDHs at DSBs are beneficial or detrimental for repair, the prevailing model suggests that early hybrid formation protects DNA ends but that excessive RDH accumulation over time hinders repair by interfering with both homologous recombination (HR) and non-homologous end joining (NHEJ).

Our data support a model where the antagonistic relationship between Nej1 and Dna2, key factors in NHEJ and HR respectively, regulates the transient accumulation of RDHs at DSBs. Although the lethality of *dna2*Δ can be supressed in combination with *pif1*-m2, the Pif1 helicase unwinds RDHs, which convinced us to utilize nuclease-deficient *dna2*-1 to investigate a role for Dna2 nuclease in hybrid resolution [40, 41, 45, 46]. In the absence of Nej1, hybrids levels reduced to background and the recovery of Dna2 increased fivefold within 45 minutes post-DSB induction, the earliest timepoint observed. In n*ej1*Δ mutants, hybrid reduction was dependent on the nuclease activity of Dna2. Nuclease-deficient *dna2*-1 not only reversed RDH loss in n*ej1*Δ, but it also increased hybrid accumulation in general, when recruited at wild type levels in *NEJ1*+ cells [15].

The impact of Dna2 on hybrid metabolism was independent of RNase H. We observed an additive increase in RDHs when *dna2*-1 was combined with *rnh201*Δ which was most prominent 90 minutes after break induction. The relative difference in hybrid levels between wild type and cells compromised in RNaseH activity by *rnh201*Δ was less pronounced at later time points, aligning with earlier work [31]. Furthermore, Dna2 overexpression significantly reduced RDHs in *rnh201*Δ mutants. Two observations strongly suggested that the role of Dna2 in hybrid resolution was not shared with Exo1 nuclease, although these two nucleases are functionally redundant in 5’ DNA resection at DSBs. First, we observed no additive increase in hybrids when *rnh201*Δ was combined with *exo1*Δ, and secondly Exo1 overexpression showed no reduction in hybrid levels in *rnh201*Δ or wild type cells even though resection increased.

Determining the effect of hybrids on resection remains a complex subject area in part because results heretofore obtained involve various experimental conditions and timepoints when hybrids and resection are measured [32]. In our experimental conditions, Dna2 is recruited, and starts resolving hybrids before Sen1 localization (Figure S6). Taken together, one model supported by our data and others is that Dna2 localization, limited by the presence of Nej1, targets hybrid metabolism early after break formation, and before it functions directly in 5’ resection (Figure 5). Thus, the presence of Nej1 enables the transient accumulation of hybrids because it limits Dna2 localization. If hybrids persist and Ku is present, then the barrier to resection increases, requiring Sen1. This model is supported by negative genetic interactions reported between *dna2* and *sen1*-1 alleles [47], and with our data wherein increased RDH levels inhibit resection independently of Ku prior to 90 minutes, but at later timepoints becomes Ku-dependent.

**Figure 5.**
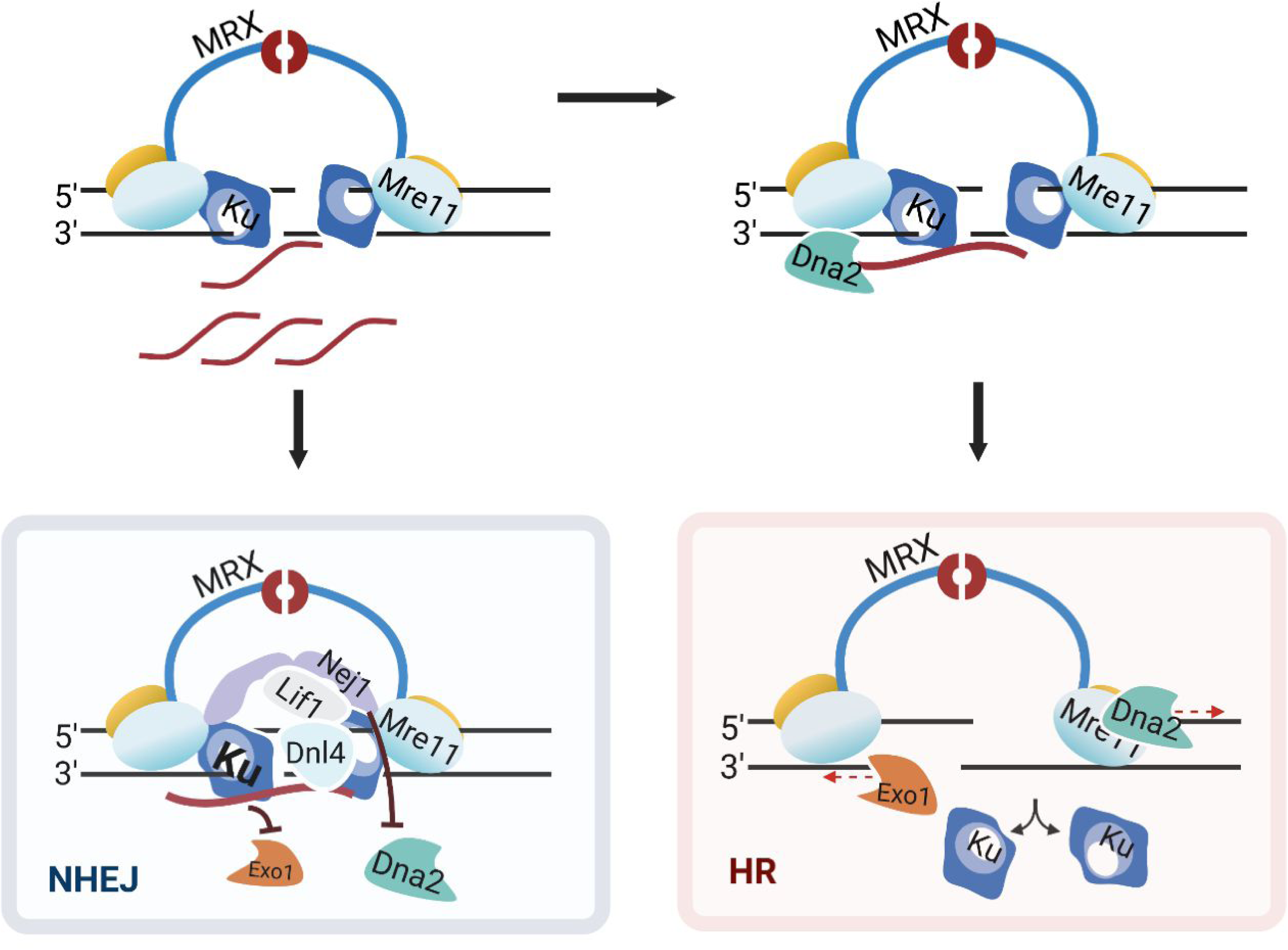
Model depicting the role and regulation of RNA:DNA hybrids in DSB repair. Upon DSB, MRX and KU complex binds the broken ends and simultaneously the pre-existing RNA (shown as red strands) transcribed from the region anneals to the broken DNA ends producing RDHs at DSB site. These hybrids enhance Ku stability and increased end-ligation (NHEJ, light blue box). As Dna2 localizes, it degrades the RDHs by targeting first duplexes with a 5’ RNA overhangs. This allows 5’ DNA resection initiation and KU disassociation for DSB repair by HR (light pink box).

Nej1 was previously shown to inhibit 5’ resection mediated by Dna2 [37], the data presented here show that Dna2-mediated resection involves multiple mechanisms. Nej1 negatively regulates the level of Dna2 recruited to the DSB, affecting both its role in long-range 5’ DNA resection and its ability to resolve RDHs at the break. The *in vitro* work showed increased Dna2 nuclease activity when RDHs with a 5’ RNA overhang was the substrate compared to substrates with a 5’ DNA overhang. Given Dna2 can initiate resection in the presence of Ku binding *in vivo*, the presence of RDHs may sequester Dna2 away from 5’ DNA temporarily. Such a mechanism would be consistent with there being a transient accumulation of hybrids at the DSB, the function of which would be to allow time for the cell to integrate DNA damage repair signals with cell cycle phase, before resection initiates, limiting options in repair pathway choice.

Given the interconnected nature of Dna2 processes affecting DNA resection, it has been challenging to pinpoint which functions directly or indirectly influence resection through RDH degradation. The observation of a mild yet reproducible improvement in the resection defect in *dna2*-1 mutants due to RnaseH1 overexpression has facilitated a clearer understanding of Dna2’s complex role in resection of DNA at DSBs. Our findings reveal how DSB repair factors regulate RDHs, adding a layer of control to DNA resection and repair pathway choice. This study elucidates the intricate interactions between RDHs and DNA repair factors, demonstrating that hybrids can significantly impact repair pathway selection by modulating resection. The research highlights the critical role of the Nej1-Dna2 interaction in governing RDHs and provides a deeper understanding of the molecular mechanisms driving DSB repair and suggest potential targets for modulating repair pathways in therapeutic applications.

## STAR ★ METHODS

### KEY RESOURCE TABLE

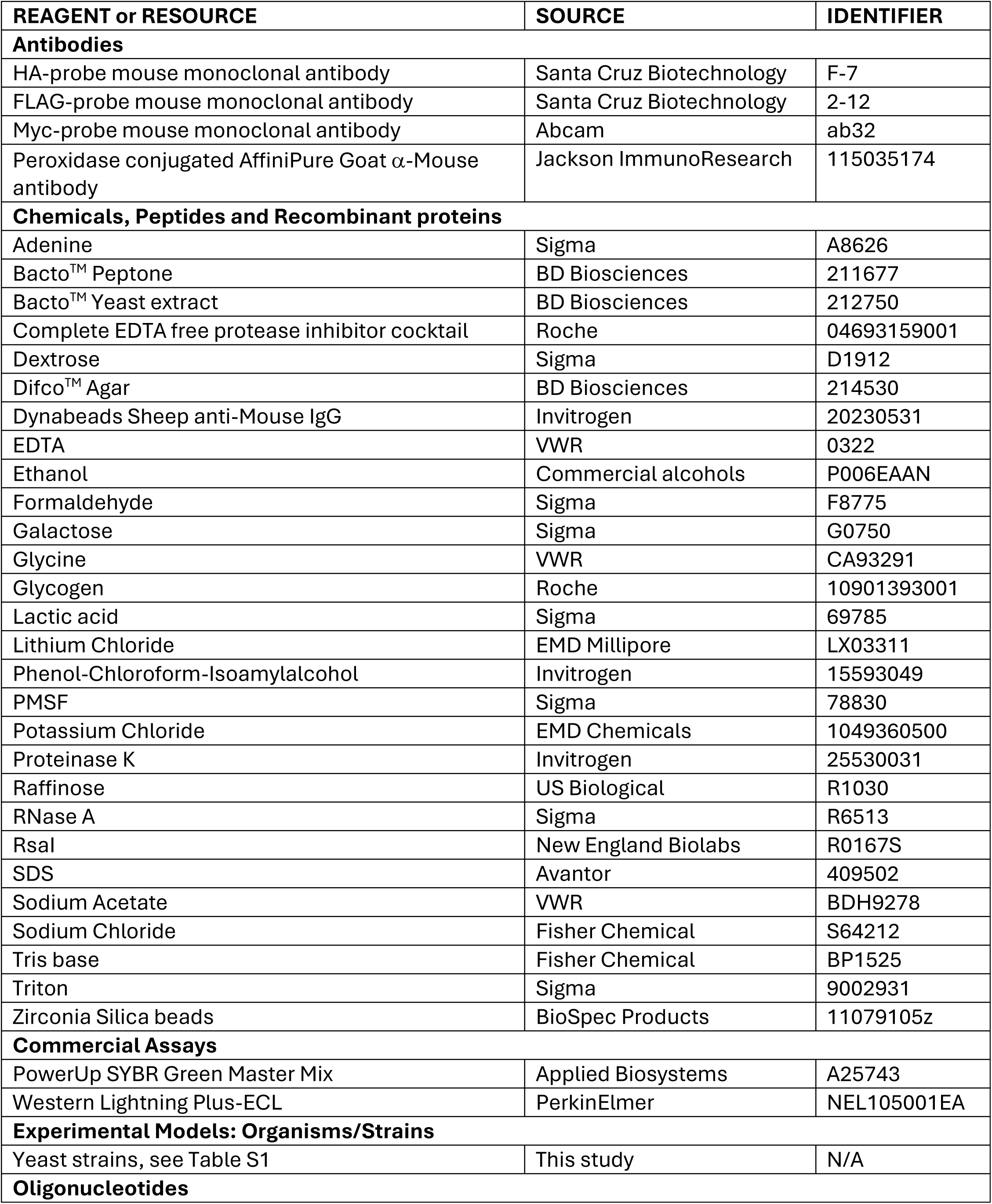
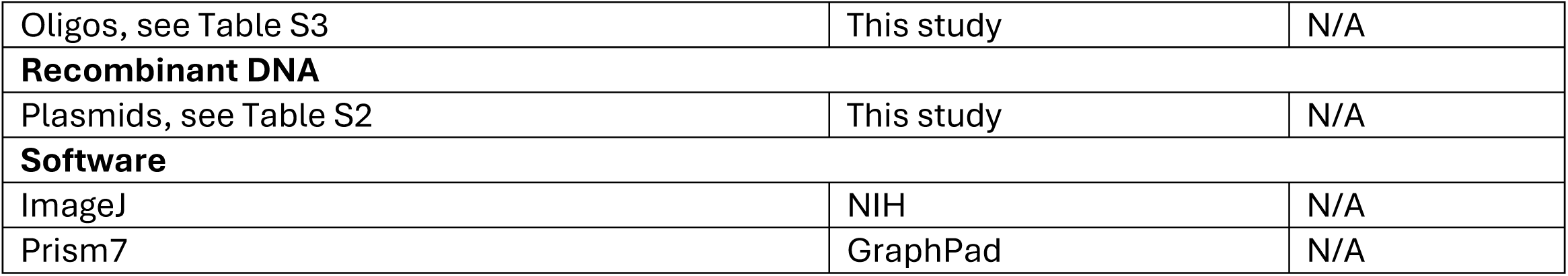

### EXPERIMENTAL MODEL AND SUBJECT DETAILS

All the yeast strains used in this study are listed in Table S1 in supplemental information and were obtained by crosses. The strains were grown on various media for the experiments and described below. For experiments involving the induction of an HO DSB, YPLG media is used (1% yeast extract, 2% bactopeptone, 2% lactic acid, 3% glycerol and 0.05% glucose). Details of plasmids and primers used in this study are specified in Table S2 and S3 in the supplemental information.

### METHOD DETAILS

#### DNA:RNA hybrid Immunoprecipitation (DRIP)

Cells were cultured overnight in YPLG at 25°C. Cells were then diluted to equal levels (5 x 10^6^ cells/ml) and were cultured to one doubling (3 hrs) at 30°C. 2% GAL was added to the YPLG and cells were harvested and crosslinked at various time points using 3.7% formaldehyde solution. Following crosslinking, the cells were washed with ice cold PBS and the pellet stored at −80°C. The pellet was re-suspended in lysis buffer (50mM Hepes pH 7.5, 1mM EDTA, 80mM NaCl, 1% Triton, 1mM PMSF and protease inhibitor cocktail) and cells were lysed using Zirconia beads and a bead beater. The lysate was sonicated to yield short DNA fragments (∼500bps in length). The sonicated lysate was then incubated in beads + S9.6 Antibody or unconjugated beads (control) for 2 hrs at 4°C. The beads were washed using lysis buffer and antibody-hybrid complex was eluted by reverse crosslinking using 1%SDS in TE buffer, followed by proteinase K treatment and nucleic acid isolation via phenol-chloroform-isoamyl alcohol extraction. Quantitative PCR was performed using the Applied Biosystem QuantStudio 6 Pro machine. PowerUp SYBR Green Master Mix was used to visualize enrichment at Mat1 (0.15kb from DSB) and PRE1 was used as an internal control.

#### Chromatin Immunoprecipitation (ChIP)

ChIP assay was performed as described previously [37]. Cells were cultured overnight in YPLG at 25°C. Cells were then diluted to equal levels (5 x 106 cells/ml) and were cultured to one doubling (3-4 hrs) at 30°C. 2% GAL was added to the YPLG and cells were harvested and crosslinked at various time points using 3.7% formaldehyde solution. Following crosslinking, the cells were washed with ice cold PBS and the pellet stored at −80°C. The pellet was re-suspended in lysis buffer (50mM Hepes pH 7.5, 1mM EDTA, 80mM NaCl, 1% Triton, 1mM PMSF and protease inhibitor cocktail) and cells were lysed using Zirconia beads and a bead beater. The lysate was sonicated to yield DNA fragments (∼500bps in length). The sonicated lysate was then incubated in beads + anti-HA/Myc/FLAG Antibody or unconjugated beads (control) for 2 hrs at 4°C. The beads were washed using wash buffer (100mM Tris pH 8, 250mM LiCl, 150mM (HA/Flag Ab) or 500mM (Myc Ab) NaCl, 0.5% NP-40, 1mM EDTA, 1mM PMSF and protease inhibitor cocktail) and protein-DNA complex was eluted by reverse crosslinking using 1%SDS in TE buffer, followed by proteinase K treatment and DNA isolation via phenol-chloroform-isoamyl alcohol extraction. Quantitative PCR was performed using the Applied Biosystem QuantStudio 6 Pro machine. PowerUp SYBR Green Master Mix was used to visualize enrichment at Mat1 (0.15kb from DSB) and PRE1 was used as an internal control.

#### qPCR based Ligation Assay

As described previously [14], cells from each strain were grown overnight in 15ml YPLG to reach an exponentially growing culture of 1×10^7^ cells/ml. Next, 2.5mL of the cells were pelleted as ‘No break’ sample, and 2% GAL was added to the remaining cells, to induce a DSB. 2.5ml of cells were pelleted after 180 minutes of incubation as timepoint 0 sample. Following that, GAL was washed off and the cells were released in YPAD and respective timepoint samples were collected. Genomic DNA was purified using standard genomic preparation method by isopropanol precipitation and ethanol washing, and DNA was re-suspended in 100uL ddH2O. Quantitative PCR was performed using the Applied Biosystem QuantStudio 6 Flex machine. PowerUp SYBR Green Master Mix was used to quantify resection at HO6 (at DSB) locus. Pre1 was used as an internal gene control for normalization. Signals from the HO6/Pre1 timepoints were normalized to ‘No break’ signals and % Ligation was determined.

#### qPCR based Resection Assay

Cells from each strain were grown overnight in 15ml YPLG to reach an exponentially growing culture of 1×10^7^ cells/ml. Next, 2.5ml of the cells were pelleted as timepoint 0 sample, and 2% GAL was added to the remaining cells, to induce a DSB. Following that, respective timepoint samples were collected. Genomic DNA was purified using standard genomic preparation method by isopropanol precipitation and ethanol washing, and DNA was re-suspended in 100mL ddH2O. Genomic DNA was treated with 0.005μg/μL RNase A for 45min at 37°C. 2μL of DNA was added to tubes containing CutSmart buffer with or without RsaI restriction enzyme and incubated at 37°C for 2hrs. Quantitative PCR was performed using the Applied Biosystem QuantStudio 6 Flex machine. PowerUp SYBR Green Master Mix was used to quantify resection at MAT1 (0.15kb from DSB) locus. Pre1 was used as a negative control. RsaI cut DNA was normalized to uncut DNA as previously described to quantify the %ssDNA [36].

#### Protein Expression and Purification

The plasmids containing the gene used for protein expression are listed in Table S2. The plasmid was transformed in wild type yeast cells (JC-727), they were grown overnight at 30°C in -TRP 2% GLU liquid media. Following day the cells were pelleted and washed thrice with autoclaved ddH2O and released in -TRP 2% GAL liquid media. The wild-type Dna2 was expressed under GAL at 25°C for 24 hrs. As done previously [14], the cells were harvested and lysed in lysis buffer (50mM Hepes pH 7.5, 1mM EDTA, 80mM NaCl, 1% Triton, 1mM PMSF, protease inhibitor cocktail and DNaseI) using Zirconia beads and a bead beater. The lysate was then incubated with beads conjugated with anti-HA Antibody for 2 hrs at 4°C. The beads were washed using wash buffer (100mM Tris pH 8, 250mM LiCl, 150mM NaCl, 0.5% NP-40, 1mM EDTA, 1mM PMSF and protease inhibitor cocktail) and protein was eluted using 100mM Glycine pH 2.0 and the eluted protein was immediately collected in 10 X TBS buffer with 60mM MgCl_2_ to avoid aggregation. Protein concentration was determined by measuring the absorbance at 280 nm, and protein purity was analyzed on SDS-PAGE and WB.

#### ThermoFluor Assay

To determine the protein stability and to compare the various mutants, a heat denaturation analysis was carried out using a CFX96 Touch real time quantitative PCR system (Bio-Rad) [48]. Protein stability measurements were performed in a buffer containing 20 mm Tris, pH 8.0, 250 mm NaCl, 5% glycerol (v/v), and 5 mm β-mercaptoethanol. The 25 μl of reaction mix contained 5 μm protein and SYPRO Orange (Invitrogen). Heat denaturing curves were observed within the temperature range of 37–85 °C, at a ramp rate of 1.8 °C/min, collecting data every 10 s. The data obtained was normalized using GraphPad Prism software.

#### Nuclease assay

Experiments were performed in a 40 μL volume in 20 mM Tris-HCl, 125 mM NaCl, 6 mM MgCl2, 1.3 mM ATP, 0.2 mg/mL BSA, 2% glycerol, pH 7.5. Reactions containing 20nM of FAM labeled probes and final protein concentrations ranging from 0 to 200nM were incubated for 30 min at 30 °C and stopped by adding 0.5% SDS, 20 mM EDTA and boiling at 95 °C for 10 min. Reactions were analyzed by running them on 20% PAGE followed by scanning the gels in Biorad Chemidoc. The bands were quantified by ImageJ software and the numbers were plotted.

#### Western Blot

Cells were lysed by re-suspending them in lysis buffer (with PMSF and protease inhibitor cocktail tablets) followed by bead beating. The protein concentration of the whole cell extract was determined using the NanoDrop. Equal amounts of whole cell extract were added to wells of 10% polyacrylamide SDS gel. After the run, the protein were transferred to Nitrocellulose membrane at 100V for 80mins. The membrane was Ponceau stained (which served as a loading control), followed by blocking in 10% milk-PBST for 1hour at room temperature. Respective primary antibody solution (1:1000 dilution) was added and left for overnight incubation at 4°C. The membranes were then washed with PBST and left for 1 hour with secondary antibody (1:10000). Followed by washing the membranes, adding the ECL substrates and imaging them.

## AUTHORS CONTRIBUTIONS

A.M. and J.A.C. designed the research. A.M. and C.G. performed experiments and analyzed the data. A.M. and J.A.C wrote the manuscript.

## ACKNOWLEDGEMENTS

This work was supported by operating grants from CIHR MOP-82736; MOP-137062 and NSERC 418122 awarded to J.A.C. The authors declare that they have no conflicts of interest with the contents of this article.

## Supplemental Information

**Figure S1.**
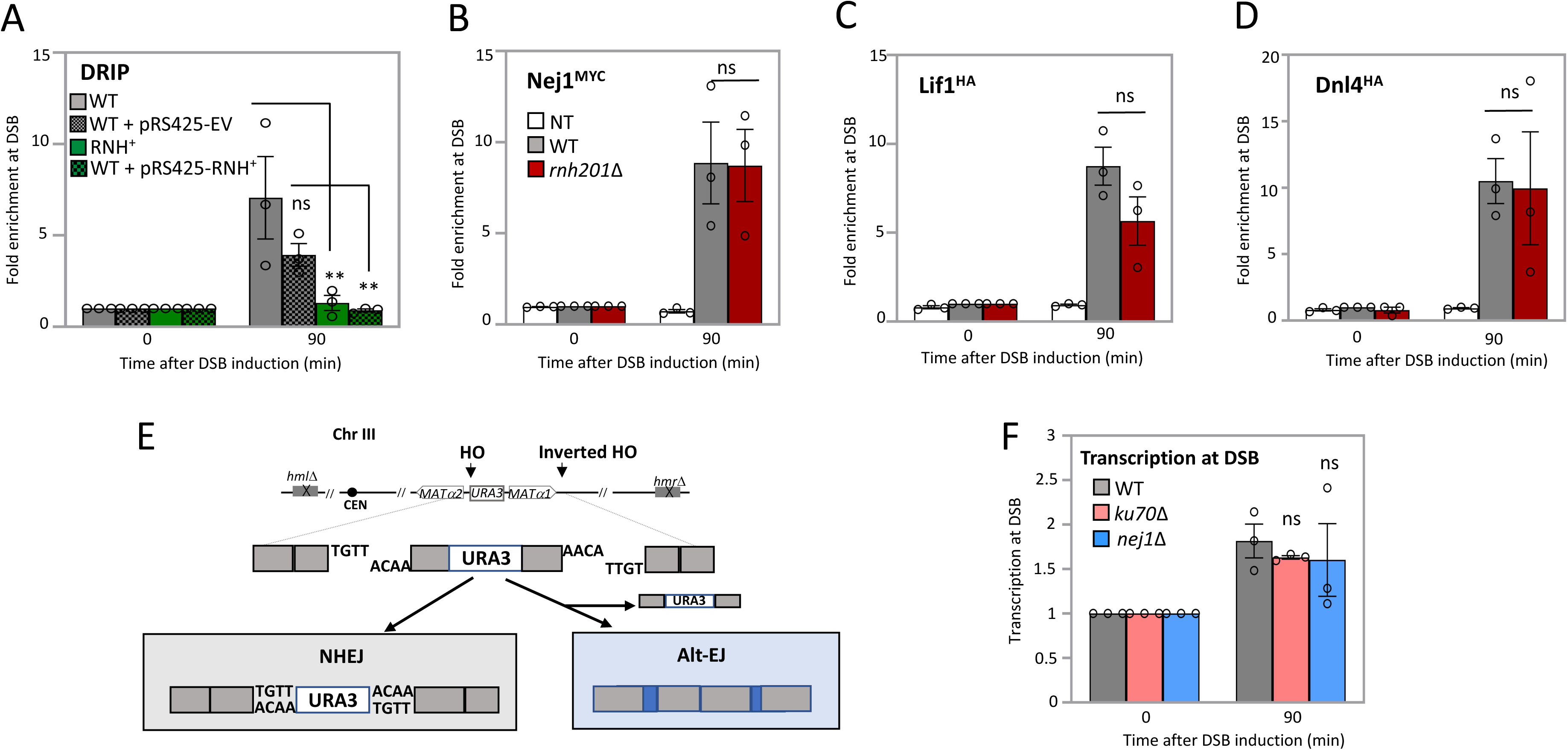
**(A)** Enrichment of RDHs using S9.6 antibody at 0.15kb from DSB, 90 mins after DSB induction relative to 0 min (no DSB induction), in wild type (JC-727) either with empty vector pRS425, or with pRS425-RnaseH1 (J-1879) and RnaseH1 *in vitro* treatment. The fold enrichment is normalized to recovery at the *PRE1* locus. **(B-D)** Enrichment of Nej1^Myc^/ Lif1^HA^/ Dnl4^HA^ at 0.15kb from DSB, 90 mins after DSB induction relative to 0 min (no DSB induction), in wild type (JC-1687/ JC-3319/ JC-5672), *rnh201*Δ (JC-5716/ JC-5715/ JC-5952), and no tag control (JC-727) was determined. The fold enrichment is normalized to recovery at the *PRE1* locus. **(E)** Schematic representation of regions around the two HO cut sites on chromosome III. Following HO endonuclease break induction, NHEJ repair joins the complementary ends of the *URA3* gene with the other ends of the break site, therefore keeping the functional *URA3* gene. From two simultaneous cuts, noncomplementary ends are generated, if repair progresses through AltEJ/MMEJ repair the URA3 gene is deleted. **(F)** Transcription at 0.15kb from DSB, 90 mins after DSB induction relative to 0 min (no DSB induction), in wild type (JC-727), *ku70*Δ (JC-1904), and *nej1*Δ (JC-1342) was determined and normalized to the *ACT1* locus. For all the experiments - error bars represent the standard error of three replicates. Significance was determined using a 1-tailed, unpaired Student’s t test (*p* < 0.05∗; *p* < 0.01∗∗; *p* < 0.001∗∗∗). All strains compared are marked and compared to WT.

**Figure S2.**
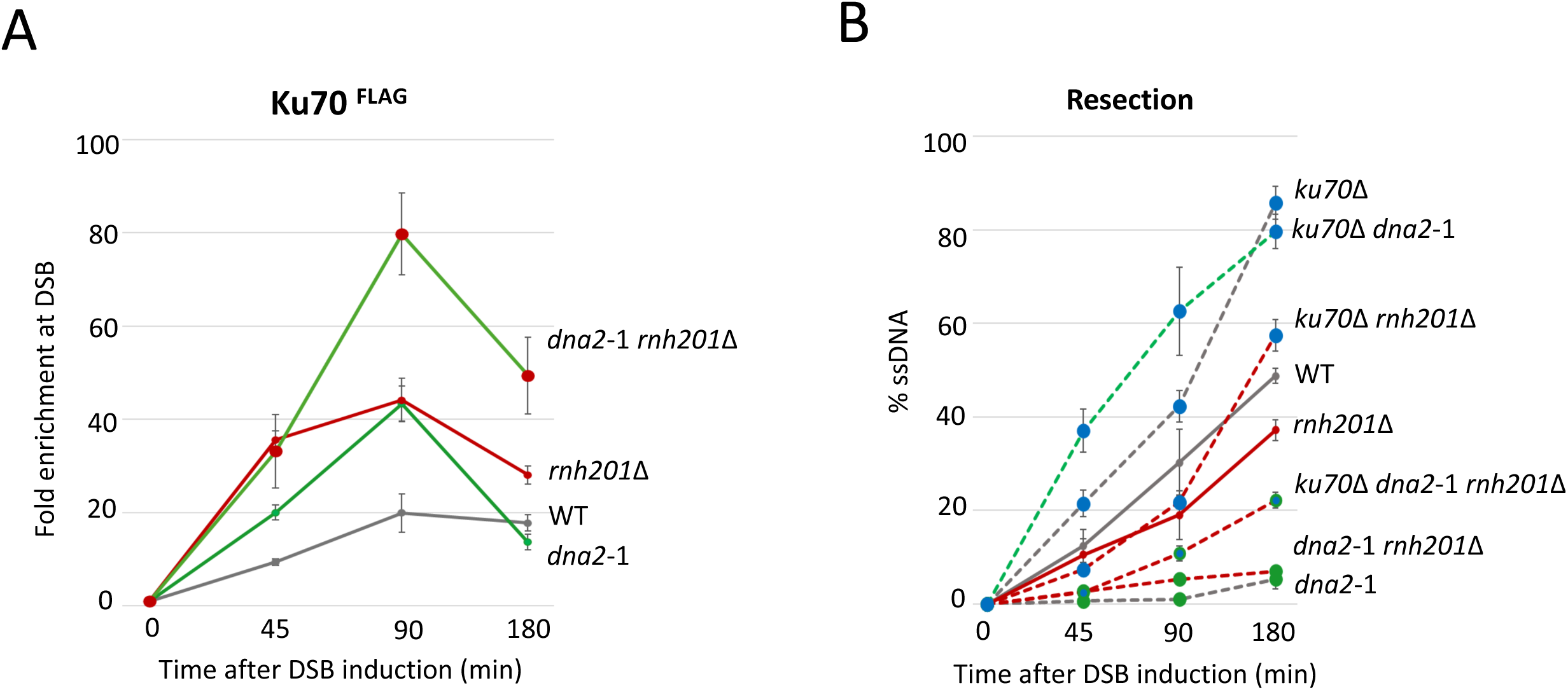
**(A)** Enrichment of Ku70^Flag^ at 0.15kb from DSB, at 0, 45, 90 and 180 mins post DSB induction in cycling cells in wild type (JC-3964), *rnh201*Δ (JC-5950), *dna2-*1 (JC-6237), and *dna2-*1 *rnh201*Δ (JC-6437) was determined. The fold enrichment is normalized to recovery at the *PRE1* locus. **(B)** qPCR-based resection assay of DNA 0.15 kb away from the HO-DSB, as measured by % ssDNA, at 0, 45, 90 and 180 mins post DSB induction in cycling cells in wild type (JC-727), *rnh201*Δ (JC-5614), *dna2-*1 (JC-6007), *dna2-*1 *rnh201*Δ (JC-5734), *ku70*Δ (JC-1904), *ku70*Δ *dna2-*1 (JC-5942), *ku70*Δ *rnh201*Δ (JC-5720), and *ku70*Δ *dna2-*1 *rnh201*Δ (JC-6441).

**Figure S3.**
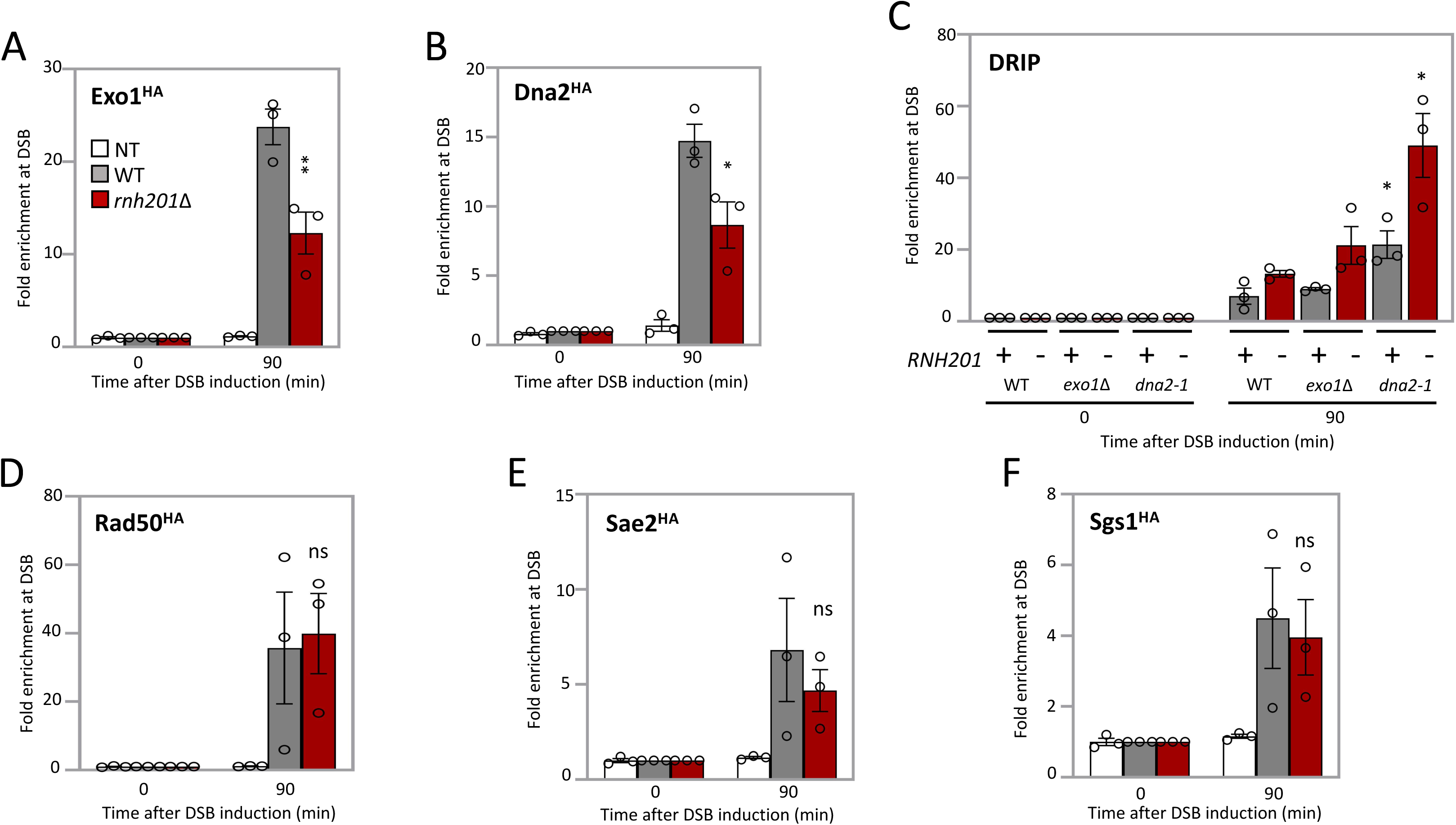
**(A-B)** Enrichment of Exo1^HA^ / Dna2^HA^ at 0.15kb from DSB, 90 mins after DSB induction relative to 0 min (no DSB induction), in wild type (JC-4869 / JC-4117), *rnh201*Δ (JC-5954 / JC-5849), and no tag control (JC-727) was determined. The fold enrichment is normalized to recovery at the *PRE1* locus. **(C)** Enrichment of RDHs using S9.6 antibody at 0.15kb from DSB, 90 mins after DSB induction relative to 0 min (no DSB induction), in wild type (JC-727), *rnh201*Δ (JC-5614), *exo1*Δ (JC-3767), *exo1*Δ *rnh201*Δ (JC-6146), *dna2-*1 (JC-6007) and *dna2-*1 *rnh201*Δ (JC-5734) was determined. The fold enrichment is normalized to recovery at the *PRE1* locus. **(D-F)** Enrichment of Rad50^HA^, Sae2^HA^, Sgs1^HA^ at 0.15kb from DSB, 90 mins after DSB induction relative to 0 min (no DSB induction), in otherwise wild type (JC-3306, JC-5116, JC-4135) or *rnh201*Δ (JC-5713, JC-5850, JC-5845), and no tag control (JC-727) was determined. The fold enrichment is normalized to recovery at the *PRE1* locus. For all the experiments - error bars represent the standard error of three replicates. Significance was determined using a 1-tailed, unpaired Student’s t test (*p* < 0.05∗; *p* < 0.01∗∗; *p* < 0.001∗∗∗). All strains compared are marked and compared to WT.

**Figure S4.**
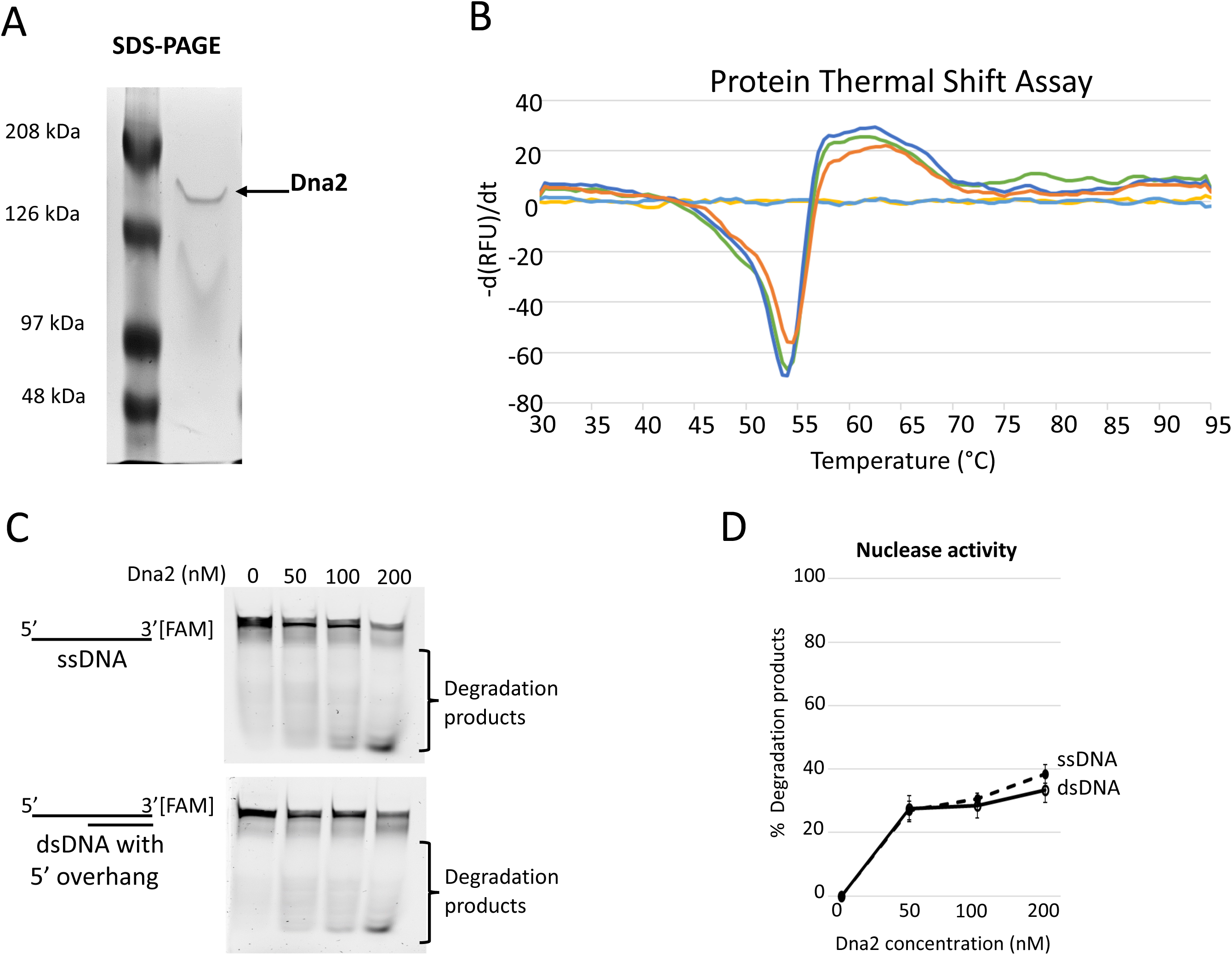
**(A)** SDS-PAGE gel of purified Dna2 protein. **(B)** Heat denaturation profiles of purified Dna2. The curves depicting the unfolded proteins in terms of −(dRFU)/dT. **(C)** Gel images showing the nuclease activity of purified Dna2 (0 to 200 nM) with 50 nM of substrates – ssDNA and dsDNA, with 5’ DNA overhang. **(D)** Plot of the quantified bands corresponding to the degradation products shown in (C) normalized to the total substrate and represented as % degradation products. Error bars represent the standard error of three replicates.

**Figure S5.**
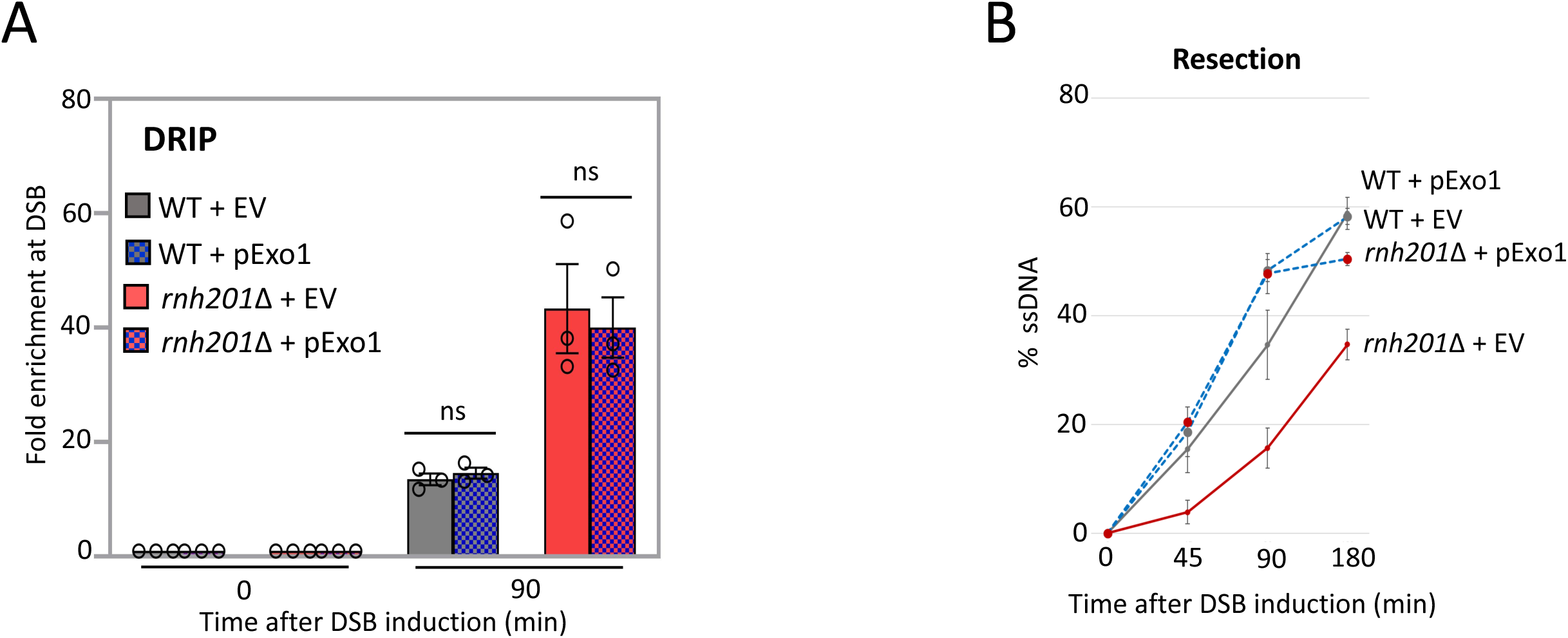
**(A)** Enrichment of RDHs using S9.6 antibody at 0.15kb from DSB, 90 mins after DSB induction relative to 0 min (no DSB induction), in wild type (JC-727), and *rnh201*Δ (JC-5614), with either 2-micron empty vector (pRS426) or 2-micron plasmid encoding Exo1 (pEM-EXO1) was determined. The fold enrichment is normalized to recovery at the *PRE1* locus. **(B)** qPCR-based resection assay of DNA 0.15 kb away from the HO-DSB, as measured by % ssDNA, at 0, 45, 90 and 180 mins post DSB induction in cycling cells in wild type (JC-727), and *rnh201*Δ (JC-5614), with either 2-micron empty vector (pRS426) or 2-micron plasmid encoding Exo1 (pEM-EXO1).

**Figure S6.**
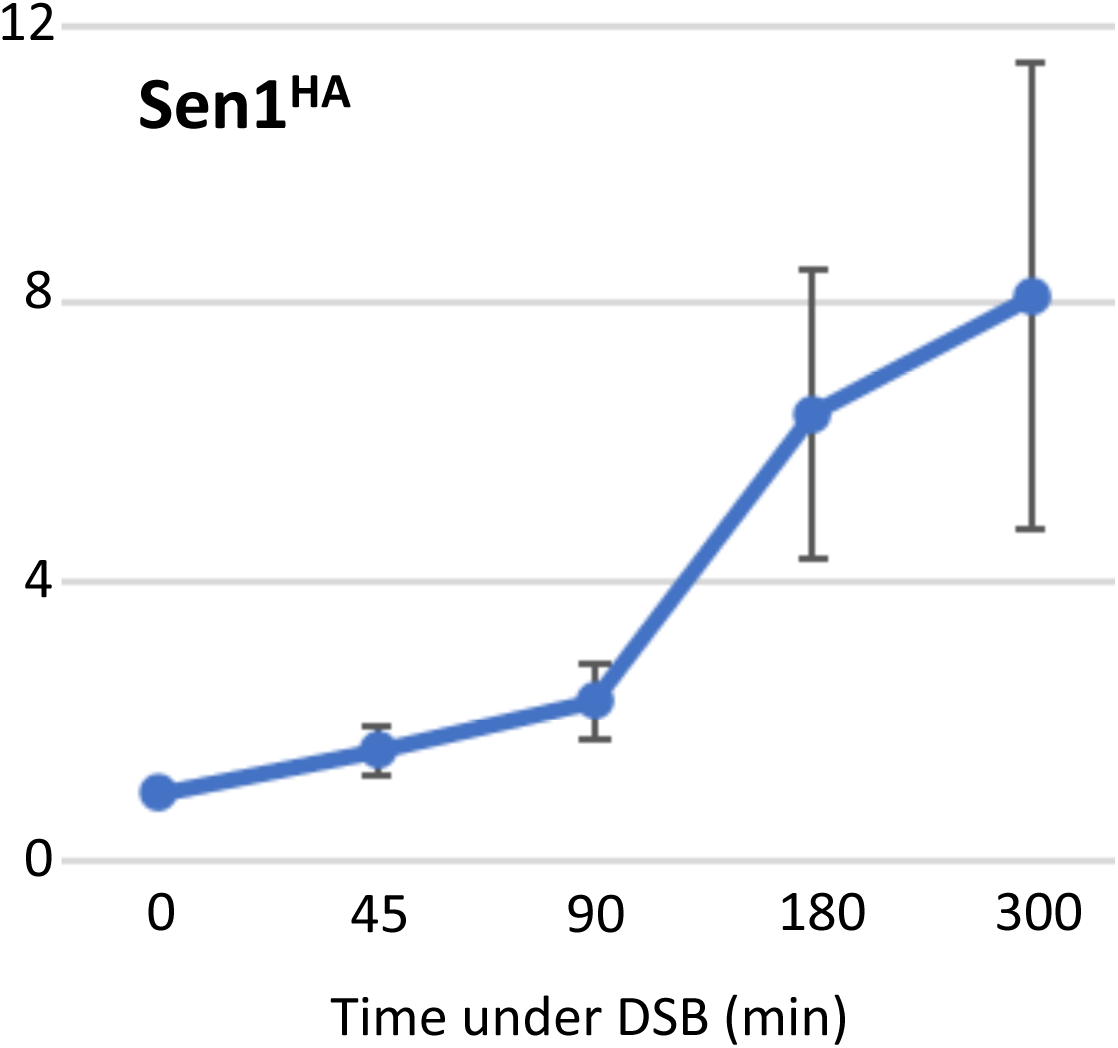
Enrichment of Sen1^HA^ (JC-6310) at 0.15kb from DSB, at 0, 45, 90, 180 and 300 mins post DSB induction in cycling cells was determined. The fold enrichment is normalized to recovery at the *PRE1* locus. Error bars represent the standard error of three replicates.

**Figure S7.**
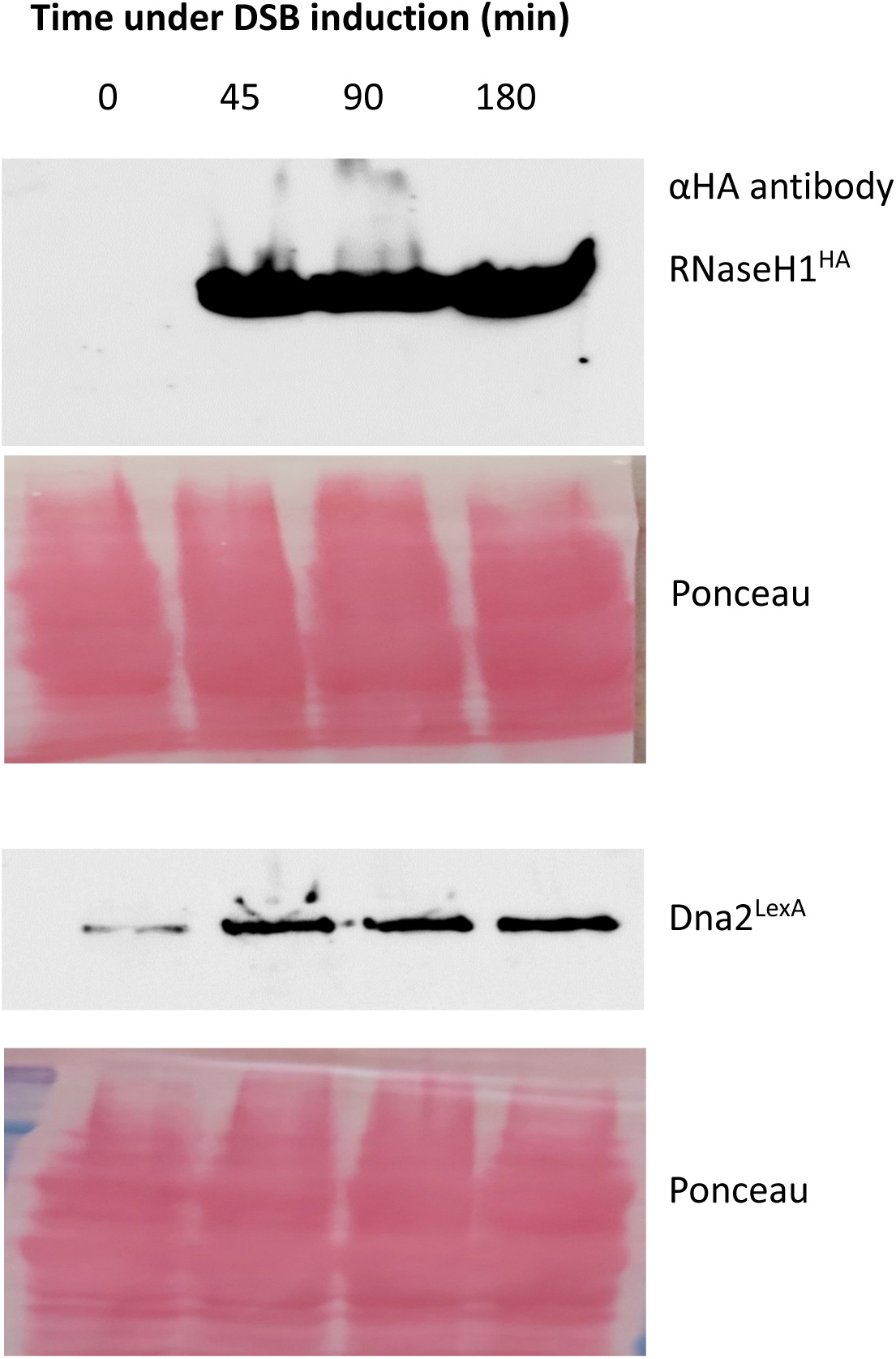
Western blot images to test RnaseH1 expression from pRS425-RnaseH1 plasmid and Dna2 expression from pGAL LexA-Dna2 plasmid, at 0, 45, 90 and 180 mins post DSB induction in cycling cells.

**Table S1.** Yeast Strains: The yeast strains used in this study are outlined.

| Strain | Genotype | Reference |
| --- | --- | --- |
| JC-727 | MAT $\alpha$; *hml::ADE1 hmr::ADE1 ade3::GAL-HO ade1-100 leu2-3, 112 lys5 trp1::hisG ura3-52* | JKM179, [Lee et al. 1998] |
| JC-1342 | JC-727 with *nej1Δ::KanMX6* | MAV015, [Valencia et al. 2001] |
| JC-1687 | JC-727 with *NEJ1-13MYC::TRP1* | Sorenson et al. 2017 |
| JC-1904 | JC-727 with *ku70Δ::KanMX6* | Sorenson et al. 2017 |
| JC-3306 | JC-727 with *RAD50-6HA::TRP1* | Sorenson et al. 2017 |
| JC-3319 | JC-727 with *LIF1-6HA::TRP1* | Sorenson et al. 2017 |
| JC-3585 | MAT $\alpha$; *hml::ADE1 hmr::ADE1 ade3::GAL-HO ade1-100 leu2-3, 112 lys5 trp1::hisG ura3-52* | JKM179, [Lee et al. 1998] |
| JC-3767 | JC-727 with *exo1Δ::NatRMX4* | Sorenson et al. 2017 |
| JC-3964 | JC-727 with *KU70-FLAG::KanMX6* | This study |
| JC-4117 | JC-727 with *DNA2-6HA::TRP1* | Mojumdar et al. 2019 |
| JC-4118 | JC-4117 with *nej1Δ::KanMX6* | Mojumdar et al. 2019 |
| JC-4135 | JC-727 with *SGS1-6HA::TRP1* | Mojumdar et al. 2019 |
| JC-4869 | JC-727 with *EXO1-6HA::TRP1* | Mojumdar et al. 2022 |
| JC-5116 | JC-727 with *SAE2-6HA::TRP1* | Mojumdar et al. 2022 |
| JC-5614 | JC-727 with *rnh201Δ::KanMX6* | This study |
| JC-5670 | JC-1342 with *dna2-1::TS* | This study |
| JC-5672 | JC-727 with *DNL4-6HA::TRP1* | Mojumdar et al. 2022 |
| JC-5713 | JC-3306 with *rnh201Δ::KanMX6* | This study |
| JC-5715 | JC-3319 with *rnh201Δ::KanMX6* | This study |
| JC-5716 | JC-1687 with *rnh201Δ::KanMX6* | This study |
| JC-5720 | JC-1904 with *rnh201Δ::KanMX6* | This study |
| JC-5733 | JC-5670 with *rnh201Δ::KanMX6* | This study |
| JC-5734 | JC-5614 with *dna2-1::TS* | This study |
| JC-5756 | JC-1342 with *rnh201Δ::KanMX6* | This study |
| JC-5845 | JC-4135 with *rnh201Δ::KanMX6* | This study |
| JC-5849 | JC-4117 with *rnh201Δ::KanMX6* | This study |
| JC-5850 | JC-5116 with *rnh201Δ::KanMX6* | This study |
| JC-5903 | MAT $\alpha$; *ho MATαpha::URA3::HOcs hml::ADE1 hmr::ADE1 ade3::GAL-HO* | Ma et al. 2003 |
| JC-5942 | JC-1904 with *dna2-1::TS* | This study |
| JC-5950 | JC-3964 with *rnh201Δ::KanMX6* | This study |
| JC-5952 | JC-5672 with *rnh201Δ::KanMX6* | This study |
| JC-5954 | JC-4869 with *rnh201Δ::KanMX6* | This study |
| JC-6007 | JC-727 with *dna2-1::TS* | Mojumdar et al. 2022 |
| JC-6146 | JC-3767 with *rnh201Δ::KanMX6* | This study |
| JC-6195 | JC-5903 with *ku70Δ::KanMX6* | This study |
| JC-6198 | JC-5903 with *rnh201Δ::KanMX6* | This study |
| JC-6237 | JC-3964 with *dna2-1::TS* | This study |
| JC-6288 | JC-6195 with *rnh201Δ::KanMX6* | This study |
| JC-6310 | JC-727 with *SEN1-3HA::TRP1* | This study |
| JC-6437 | JC-5950 with *dna2-1::TS* | This study |
| JC-6441 | JC-5720 with *dna2-1::TS* | This study |

**Table S2.** Plasmids: Table of Plasmids used in this study.

| Plasmid Number | Description |
| --- | --- |
| J-282 | pRS426-Empty vector |
| J-965 | pGAL LexA-Empty vector |
| J-1853 | pGAL LexA-Dna2 |
| J-1879 | pRS425-Rnh1 |
| J-1880 | pRS425-Empty vector |
| J-1952 | pRS426-Exo1 |

**Table S3.**
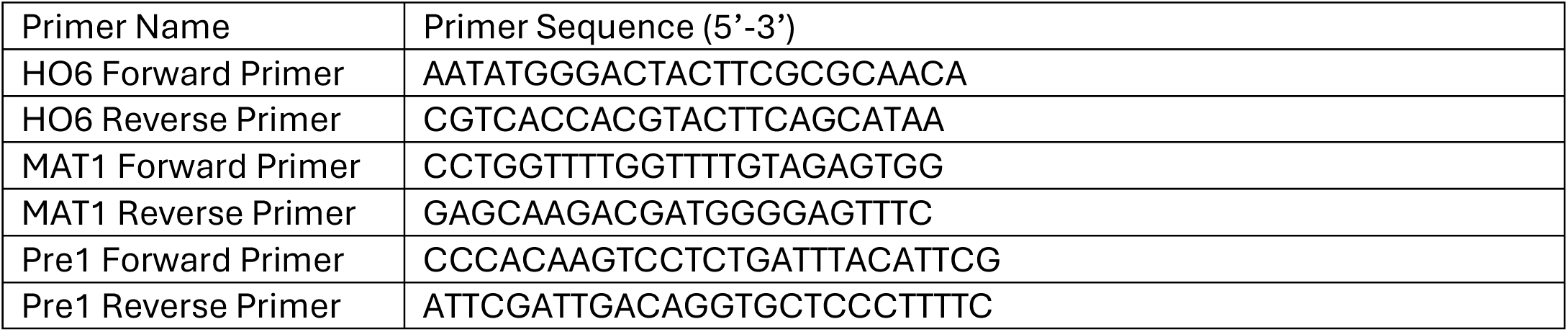
Primers: Table of Primers and Probes used in this study.

**Table S4:** Table of DSB cut efficiency for strains used in this study.

| Strain | DSB cut (%) |  |  |
| --- | --- | --- | --- |
|  | 45min | 90min | 180min |
| JC-727 | 89.11 | 95.37 | 96.72 |
| JC-1342 | 83.35 | 93.49 | 95.85 |
| JC-1904 | 95.75 | 98.20 | 98.59 |
| JC-3964 | 98.99 | 98.77 | 98.56 |
| JC-4117 | 98.76 | 98.68 | 98.51 |
| JC-4118 | 98.97 | 98.63 | 98.83 |
| JC-5614 | 80.11 | 81.15 | 88.84 |
| JC-5670 | 98.74 | 98.61 | 98.26 |
| JC-5720 | 96.97 | 97.56 | 98.64 |
| JC-5733 | 98.88 | 99.36 | 98.85 |
| JC-5734 | 84.45 | 91.81 | 95.13 |
| JC-5756 | 80.46 | 88.75 | 91.20 |
| JC-5942 | 94.84 | 98.41 | 98.50 |
| JC-5950 | 96.27 | 98.07 | 98.39 |
| JC-6007 | 88.16 | 90.50 | 95.48 |
| JC-6237 | 94.66 | 97.48 | 98.37 |
| JC-6437 | 95.99 | 98.56 | 98.87 |
| JC-6310 | 98.95 | 98.92 | 99.25 |
| JC-6441 | 96.41 | 95.26 | 98.44 |

**Table S5:** Table of DSB cut efficiency for strains used in this study.

| Strain | DSB cut % at 90min |
| --- | --- |
| JC-1687 | 95.24 |
| JC-3306 | 94.37 |
| JC-3313 | 95.21 |
| JC-3319 | 94.83 |
| JC-3767 | 94.19 |
| JC-4135 | 95.96 |
| JC-4869 | 94.66 |
| JC-5116 | 94.65 |
| JC-5672 | 96.20 |
| JC-5713 | 95.44 |
| JC-5715 | 95.62 |
| JC-5716 | 94.97 |
| JC-5845 | 94.22 |
| JC-5849 | 95.49 |
| JC-5850 | 98.46 |
| JC-5950 | 95.01 |
| JC-5952 | 98.92 |
| JC-5954 | 95.83 |
| JC-6146 | 95.55 |

## References

1. Symington, LS. (2016). Mechanism and regulation of DNA end resection in eukaryotes. Crit Rev Biochem Mol Biol. 51(3), 195–212. 10.3109/10409238.2016.1172552

2. Wu, D., Topper, L. M., and Wilson, T. E. (2008). Recruitment and dissociation of nonhomologous end joining proteins at a DNA double-strand break in Saccharomyces cerevisiae. Genetics, 178(3), 1237–1249. 10.1534/genetics.107.083535

3. Palmbos, P. L., Wu, D., Daley, J. M., and Wilson, T. E. (2008). Recruitment of Saccharomyces cerevisiae Dnl4-Lif1 complex to a double-strand break requires interactions with Yku80 and the Xrs2 FHA domain. Genetics, 180(4), 1809–1819. 10.1534/genetics.108.095539

4. Hopfner, K.-P., Craig, L., Moncalian, G., Zinkel, R. a, Usui, T., Owen, B. a L., Karcher, A., Henderson, B., Bodmer, J.-L., McMurray, C. T., et al. (2002) The Rad50 zinc-hook is a structure joining Mre11 complexes in DNA recombination and repair. Nature. 418, 562–566. 10.1038/nature00922

5. Wiltzius, J. J., Hohl, M., Fleming, J. C., and Petrini, J. H. (2005). The Rad50 hook domain is a critical determinant of Mre11 complex functions. Nature structural & molecular biology, 12(5), 403–407. 10.1038/nsmb928

6. Marini, F., Rawal, C. C., Liberi, G., and Pellicioli, A. (2019). Regulation of DNA Double Strand Breaks Processing: Focus on Barriers. Frontiers in molecular biosciences, 6, 55. 10.3389/fmolb.2019.00055

7. Cejka P. (2015). DNA End Resection: Nucleases Team Up with the Right Partners to Initiate Homologous Recombination. The Journal of biological chemistry, 290(38), 22931–22938. 10.1074/jbc.R115.675942

8. Cannavo, E., and Cejka, P. (2014). Sae2 promotes dsDNA endonuclease activity within Mre11-Rad50-Xrs2 to resect DNA breaks. Nature, 514(7520), 122–125. 10.1038/nature13771

9. Zhu, Z., Chung, W. H., Shim, E. Y., Lee, S. E., and Ira, G. (2008). Sgs1 helicase and two nucleases Dna2 and Exo1 resect DNA double-strand break ends. Cell, 134(6), 981–994. 10.1016/j.cell.2008.08.037

10. Gobbini, E., Villa, M., Gnugnoli, M., Menin, L., Clerici, M., and Longhese, M. P. (2015). Sae2 Function at DNA Double-Strand Breaks Is Bypassed by Dampening Tel1 or Rad53 Activity. PLoS genetics, 11(11), e1005685. 10.1371/journal.pgen.1005685

11. Yu, T. Y., Kimble, M. T., and Symington, L. S. (2018). Sae2 antagonizes Rad9 accumulation at DNA double-strand breaks to attenuate checkpoint signaling and facilitate end resection. Proceedings of the National Academy of Sciences of the United States of America, 115(51), E11961–E11969. 10.1073/pnas.1816539115

12. Arora, S., Deshpande, R. A., Budd, M., Campbell, J., Revere, A., Zhang, X., Schmidt, K. H., and Paull, T. T. (2017). Genetic Separation of Sae2 Nuclease Activity from Mre11 Nuclease Functions in Budding Yeast. Molecular and cellular biology, 37(24), e00156–17. 10.1128/MCB.00156-17

13. Mojumdar, A., Adam, N., and Cobb, J. A. (2022). Nej1 interacts with Sae2 at DNA double-stranded breaks to inhibit DNA resection. The Journal of biological chemistry, 298(6), 101937. 10.1016/j.jbc.2022.101937

14. Mojumdar, A., Adam, N., and Cobb, J. A. (2022). Multifunctional properties of Nej1XLF C-terminus promote end-joining and impact DNA double-strand break repair pathway choice. DNA repair, 115, 103332. 10.1016/j.dnarep.2022.103332

15. Mojumdar, A., Granger, C., Lunke, M., and Cobb, J. A. (2024). Loss of Dna2 fidelity results in decreased Exo1-mediated resection at DNA double-strand breaks. The Journal of biological chemistry, 300(3), 105708. 10.1016/j.jbc.2024.105708

16. Jimeno, S., Prados-Carvajal, R., and Huertas, P. (2019). The role of RNA and RNA-related proteins in the regulation of DNA double strand break repair pathway choice. DNA repair, 81, 102662. 10.1016/j.dnarep.2019.102662

17. Puget, N., Miller, K. M., and Legube, G. (2019). Non-canonical DNA/RNA structures during Transcription-Coupled Double-Strand Break Repair: Roadblocks or Bona fide repair intermediates?. DNA repair, 81, 102661. 10.1016/j.dnarep.2019.102661

18. Ohle, C., Tesorero, R., Schermann, G., Dobrev, N., Sinning, I., and Fischer, T. (2016). Transient RNA-DNA Hybrids Are Required for Efficient Double-Strand Break Repair. Cell, 167(4), 1001–1013.e7. 10.1016/j.cell.2016.10.001

19. Marnef, A., and Legube, G. (2021). R-loops as Janus-faced modulators of DNA repair. Nature cell biology, 23(4), 305–313. 10.1038/s41556-021-00663-4

20. Pessina, F., Giavazzi, F., Yin, Y., Gioia, U., Vitelli, V., Galbiati, A., Barozzi, S., Garre, M., Oldani, A., Flaus, A., et al. (2019). Functional transcription promoters at DNA double-strand breaks mediate RNA-driven phase separation of damage-response factors. Nature cell biology, 21(10), 1286–1299. 10.1038/s41556-019-0392-4.

21. Ortega, P., Mérida-Cerro, J. A., Rondón, A. G., Gómez-González, B., and Aguilera, A. (2021). DNA-RNA hybrids at DSBs interfere with repair by homologous recombination. eLife, 10, e69881. 10.7554/eLife.69881

22. Domingo-Prim, J., Bonath, F., and Visa, N. (2020). RNA at DNA Double-Strand Breaks: The Challenge of Dealing with DNA:RNA Hybrids. BioEssays : news and reviews in molecular, cellular and developmental biology, 42(5), e1900225. 10.1002/bies.201900225.

23. Michelini, F., Pitchiaya, S., Vitelli, V., Sharma, S., Gioia, U., Pessina, F., Cabrini, M., Wang, Y., Capozzo, I., Iannelli, F., et al. (2017). Damage-induced lncRNAs control the DNA damage response through interaction with DDRNAs at individual double-strand breaks. Nature cell biology, 19(12), 1400–1411. 10.1038/ncb3643.

24. Audoynaud, C., Vagner, S., and Lambert, S. (2021). Non-homologous end-joining at challenged replication forks: an RNA connection?. Trends in genetics : TIG, 37(11), 973–985. 10.1016/j.tig.2021.06.010

25. Sessa, G., Gómez-González, B., Silva, S., Pérez-Calero, C., Beaurepere, R., Barroso, S., Martineau, S., Martin, C., Ehlén, Å., Martínez, J. S., et al. (2021). BRCA2 promotes DNA-RNA hybrid resolution by DDX5 helicase at DNA breaks to facilitate their repair‡. The EMBO journal, 40(7), e106018. 10.15252/embj.2020106018.

26. D’Alessandro, G., Whelan, D. R., Howard, S. M., Vitelli, V., Renaudin, X., Adamowicz, M., Iannelli, F., Jones-Weinert, C. W., Lee, M., Matti, V., et al. (2018). BRCA2 controls DNA:RNA hybrid level at DSBs by mediating RNase H2 recruitment. Nature communications, 9(1), 5376. 10.1038/s41467-018-07799-2.

27. Li, L., Germain, D. R., Poon, H. Y., Hildebrandt, M. R., Monckton, E. A., McDonald, D., Hendzel, M. J., and Godbout, R. (2016). DEAD Box 1 Facilitates Removal of RNA and Homologous Recombination at DNA Double-Strand Breaks. Molecular and cellular biology, 36(22), 2794–2810. 10.1128/MCB.00415-16

28. Yasuhara, T., Kato, R., Hagiwara, Y., Shiotani, B., Yamauchi, M., Nakada, S., Shibata, A., and Miyagawa, K. (2018). Human Rad52 Promotes XPG-Mediated R-loop Processing to Initiate Transcription-Associated Homologous Recombination Repair. Cell, 175(2), 558–570.e11. 10.1016/j.cell.2018.08.056

29. Chakraborty, A., Tapryal, N., Venkova, T., Horikoshi, N., Pandita, R. K., Sarker, A. H., Sarkar, P. S., Pandita, T. K., and Hazra, T. K. (2016). Classical non-homologous end-joining pathway utilizes nascent RNA for error-free double-strand break repair of transcribed genes. Nature communications, 7, 13049. 10.1038/ncomms13049

30. Zong, D., Oberdoerffer, P., Batista, P. J., and Nussenzweig, A. (2020). RNA: a double-edged sword in genome maintenance. Nature reviews. Genetics, 21(11), 651–670. 10.1038/s41576-020-0263-7

31. Rawal, C. C., Zardoni, L., Di Terlizzi, M., Galati, E., Brambati, A., Lazzaro, F., Liberi, G., and Pellicioli, A. (2020). Senataxin Ortholog Sen1 Limits DNA:RNA Hybrid Accumulation at DNA Double-Strand Breaks to Control End Resection and Repair Fidelity. Cell reports, 31(5), 107603. 10.1016/j.celrep.2020.107603

32. Gómez-González, B., and Aguilera, A. (2023). Break-induced RNA-DNA hybrids (BIRDHs) in homologous recombination: friend or foe?. EMBO reports, 24(12), e57801. 10.15252/embr.202357801

33. Audoynaud, C., Schirmeisen, K., Ait Saada, A., Gesnik, A., Fernández-Varela, P., Boucherit, V., Ropars, V., Chaudhuri, A., Fréon, K., Charbonnier, J. B., and Lambert, S. A. E. (2023). RNA:DNA hybrids from Okazaki fragments contribute to establish the Ku-mediated barrier to replication-fork degradation. Molecular cell, 83(7), 1061– 1074.e6. 10.1016/j.molcel.2023.02.008

34. Haber, JE. (2012). Mating-type genes and MAT switching in Saccharomyces cerevisiae. Genetics,191(1), 33–64. 10.1534/genetics.111.134577

35. Zhao, H., Zhu, M., Limbo, O., and Russell, P. (2018). RNase H eliminates R-loops that disrupt DNA replication but is nonessential for efficient DSB repair. EMBO reports, 19(5), e45335. 10.15252/embr.201745335

36. Ferrari, M., Dibitetto, D., De Gregorio, G., Eapen, V. V., Rawal, C. C., Lazzaro, F., Tsabar, M., Marini, F., Haber, J. E., and Pellicioli, A. (2015). Functional interplay between the 53BP1-ortholog Rad9 and the Mre11 complex regulates resection, end-tethering and repair of a double-strand break. PLoS genetics, 11(1), e1004928. 10.1371/journal.pgen.1004928

37. Mojumdar, A., Sorenson, K., Hohl, M., Toulouze, M., Lees-Miller, S. P., Dubrana, K., Petrini, J. H. J., and Cobb, J. A. (2019). Nej1 Interacts with Mre11 to Regulate Tethering and Dna2 Binding at DNA Double-Strand Breaks. Cell reports, 28(6), 1564–1573.e3. 10.1016/j.celrep.2019.07.018

38. Ma, J. L., Kim, E. M., Haber, J. E., and Lee, S. E. (2003). Yeast Mre11 and Rad1 proteins define a Ku-independent mechanism to repair double-strand breaks lacking overlapping end sequences. Molecular and cellular biology, 23(23), 8820– 8828. 10.1128/MCB.23.23.8820-8828.2003

39. Mimitou, E. P., and Symington, L. S. (2010). Ku prevents Exo1 and Sgs1-dependent resection of DNA ends in the absence of a functional MRX complex or Sae2. The EMBO journal, 29(19), 3358–3369. 10.1038/emboj.2010.193

40. Budd, M. E., and Campbell, J. L. (1995). A yeast gene required for DNA replication encodes a protein with homology to DNA helicases. Proceedings of the National Academy of Sciences of the United States of America, 92(17), 7642–7646. 10.1073/pnas.92.17.7642

41. Budd, M. E., Choe, W.c, and Campbell, J. L. (2000). The nuclease activity of the yeast DNA2 protein, which is related to the RecB-like nucleases, is essential in vivo. The Journal of biological chemistry, 275(22), 16518–16529. 10.1074/jbc.M909511199

42. Jia, P. P., Junaid, M., Ma, Y. B., Ahmad, F., Jia, Y. F., Li, W. G., and Pei, D. S. (2017). Role of human DNA2 (hDNA2) as a potential target for cancer and other diseases: A systematic review. DNA repair, 59, 9–19. 10.1016/j.dnarep.2017.09.001

43. Kumar, S., Peng, X., Daley, J., Yang, L., Shen, J., Nguyen, N., Bae, G., Niu, H., Peng, Y., Hsieh, H. J., et al. (2017). Inhibition of DNA2 nuclease as a therapeutic strategy targeting replication stress in cancer cells. Oncogenesis, 6(4), e319. 10.1038/oncsis.2017.15

44. Bae, S. H., Bae, K. H., Kim, J. A., and Seo, Y. S. (2001). RPA governs endonuclease switching during processing of Okazaki fragments in eukaryotes. Nature, 412(6845), 456–461. 10.1038/35086609

45. Budd, M. E., Reis, C. C., Smith, S., Myung, K., and Campbell, J. L. (2006). Evidence suggesting that Pif1 helicase functions in DNA replication with the Dna2 helicase/nuclease and DNA polymerase delta. Molecular and cellular biology, 26(7), 2490–2500. 10.1128/MCB.26.7.2490-2500.2006

46. Boulé, J. B., and Zakian, V. A. (2007). The yeast Pif1p DNA helicase preferentially unwinds RNA DNA substrates. Nucleic acids research, 35(17), 5809–5818. 10.1093/nar/gkm613

47. Costanzo, M., VanderSluis, B., Koch, E. N., Baryshnikova, A., Pons, C., Tan, G., Wang, W., Usaj, M., Hanchard, J., Lee, S. D., et al. (2016). A global genetic interaction network maps a wiring diagram of cellular function. Science (New York, N.Y.), 353(6306), aaf1420. 10.1126/science.aaf1420

48. Mojumdar, A., De March, M., Marino, F., and Onesti, S. (2017). The Human RecQ4 Helicase Contains a Functional RecQ C-terminal Region (RQC) That Is Essential for Activity. The Journal of biological chemistry, 292(10), 4176–4184. 10.1074/jbc.M116.767954

